# Apical-Out Ileal Organoids: A Versatile Platform for Probing Gut-Nanomaterial Interactions

**DOI:** 10.64898/2026.10.10.758216

**Authors:** Dinh Son Vo, Alina Hartwig, Foteini Tzioufa, Jens Eriksson, Ana C. C. Lopes, Maria Letizia Di Martino, Wojciech Fijalkowski, James Rae, Kajsa Björner, Johan Vessby, Robert G. Parton, Hanna M. G. Barriga, Mikael E. Sellin, Per Artursson, Madlen Hubert

## Abstract

The human ileal epithelium forms a critical interface governing interactions with luminal microorganisms and nanomaterials, yet physiologically relevant in vitro models with accessible apical surfaces remain limited. Here, human apical-out ileoids derived from healthy ileal biopsies are established as a platform that recapitulates the architecture and complexity of the native ileal epithelium. Assessment of donor- and passage-dependent variability identifies a representative ileoid line and a robust culture window. Integrated proteomic, imaging, ultrastructural, and functional analyses demonstrate preservation of key epithelial features, including brush-border organization, barrier integrity, absorptive capacity, and multiple endocytic pathways. Intact ileoids internalize nanoparticles of distinct surface charge, including empty lipid nanoparticles compositionally matched to approved mRNA vaccine formulations, establishing that uptake can be quantified across an unperturbed apical surface. This uptake is restrained by a retained mucin-rich glycocalyx. Cleavage of surface-attached mucins selectively enhances uptake of negatively charged nanoparticles, leaving near-neutral and positively charged nanoparticles unchanged, and increasing entry of *Salmonella* bacteria. These findings reveal the glycocalyx as a charge-selective regulator of epithelial uptake. In conclusion, our study provides a foundation for nanoparticle-epithelium investigations using human apical-out ileoids, and a versatile, translationally relevant platform for nanomedicine delivery, host-microbe interactions, and epithelial transport.

## 1. Introduction

Reliable *in vitro* models of the human intestinal epithelium are essential for early-stage drug discovery and for advancing the 3R principles (replacement, reduction and refinement) in animal biomedical research.^[1]^ Conventional two-dimensional (2D) models remain widely used because they are relatively inexpensive, robust and easy to culture.^[2]^ Among these, the Caco-2 cell line, derived from human colon adenocarcinoma, is widely regarded as a standard model for assessing intestinal drug permeability.^[3,4]^ To improve physiological relevance, co-culture systems such as Caco-2/HT29 and Caco-2/Raji models have been developed to introduce mucus-producing goblet cells or M-cell characteristics, respectively.^[5–8]^ Despite these advances, conventional 2D models do not adequately reproduce the cellular diversity, three-dimensional architecture, crypt-villus organization, and complex epithelial interactions of the human intestine.

Intestinal organoids have emerged as physiologically relevant three-dimensional (3D) models that reproduce many structural and functional characteristics of the native intestinal epithelium, including epithelial cell diversity and tissue-specific functions.^[9–12]^ Adult stem cell-derived organoids (denoted “enteroids” or “colonoids” depending on the segment of origin) are typically established from isolated intestinal crypts embedded in extracellular matrix (ECM), where intestinal stem cells self-organize and give rise to multicellular structures that contain multiple epithelial cell types.^[9,10,12,13]^ Their compatibility with advanced imaging, genetic profiling and editing, and omics technologies has made organoids valuable tools for investigating intestinal physiology, host-microbe interactions, disease mechanisms, and drug absorption, metabolism, and toxicity.^[9,14–17]^ Still, intestinal organoids remain subject to important experimental limitations. Primary cells may undergo progressive cellular senescence and phenotypic or epigenetic change during extended culture and serial passaging, while donor-specific characteristics, including age and disease state, may influence regenerative capacity and contribute to variability between organoid lines.^[18–20]^ Systematic monitoring of organoid quality and careful selection of representative donor lines are therefore important for improving experimental reproducibility.

A second major limitation of conventional ECM-embedded intestinal organoids is their basal-out polarity, which restricts direct access to the luminal or apical, surface.^[10]^ Consequently, studies requiring controlled apical exposure, including investigations of epithelial transport, host-microbe interactions, and nanoparticle uptake, often rely on technically demanding approaches such as microinjection.^[15,21,22]^ In addition, the surrounding ECM can influence the diffusion, accessibility, and effective concentration of externally applied compounds.^[23]^ Recent studies have demonstrated that removal of the surrounding ECM can induce polarity reversal, generating apical-out organoids in which the brush border and apical membrane face the surrounding culture medium.^[9,10]^ This configuration provides direct and simultaneous access to the epithelial apical surface across the organoid population, facilitating exposure to luminal compounds, microorganisms, and drug delivery systems. Apical-out organoids therefore offer an attractive platform for studying epithelial transport and interactions at the intestinal epithelial interface, while avoiding the technical limitations associated with microinjection.

The ileum is of particular interest for such applications because it represents one of the principal sites affected in Crohn’s disease, with isolated ileal involvement occurring in approximately 30–40% of patients.^[24,25]^ Similar to duodenum and jejunum, the ileal epithelium is composed of a highly specialized and dynamic population of absorptive and secretory cells organized within a crypt-villus architecture, enabling functions ranging from nutrient and drug absorption to barrier defence and host-microbe interactions.^[26,27]^ The apical surface is further covered by a glycocalyx composed of membrane-associated mucins, proteoglycans, glycosaminoglycans, and other glycoconjugates.^[28–31]^ This carbohydrate-rich interface contributes to epithelial protection and regulates interactions with luminal microorganisms, nutrients, and therapeutic cargo.^[32]^ In the context of oral drug delivery and nanomedicine, the glycocalyx may influence the adhesion, diffusion, and cellular uptake of nanoparticles through its physicochemical properties, including steric and electrostatic interactions.^[33–36]^ Despite its potential importance, the structural and functional contribution of the human ileal glycocalyx to interactions with nanomaterials and biological particles remains incompletely understood. This is partly attributable to the limited availability of physiologically relevant human models that provide direct experimental access to the apical epithelial interface.

Although apical-out intestinal organoids provide an opportunity to overcome this limitation, their suitability as standardized models of the human ileal epithelial surface has not yet been comprehensively established.^[37]^ In particular, donor-dependent variability, passage-related changes, long-term stability, epithelial phenotype and function, and the integrity of the apical glycocalyx remain only partially explored.^[18–20]^ Establishing a robust and reproducible apical-out ileal model is therefore important not only for studying epithelial physiology and host-microbe interactions, but also for investigating how the human intestinal surface interacts with orally administered nanomaterials. This is particularly relevant for nanoparticle studies, which commonly rely on conventional 2D models such as Caco-2 cells.^[38,39]^ A physiologically relevant human model with direct access to the apical glycocalyx could provide complementary insight into nanoparticle interactions with and uptake by the intestinal epithelium

Here, we establish and comprehensively validate a standardized pipeline for generating apical-out human ileal enteroids (hereafter “ileoids”) from healthy ileal biopsies. We define experimental workflow parameters incorporating donor selection and passage optimization, characterize the resulting model at structural, molecular, and functional levels, and establish the presence and functional relevance of the apical ileal glycocalyx. We further investigate how disruption of surface-associated mucins in the glycocalyx influences the interaction between the epithelial surface and biological particles. To this end, a panel of nanoparticles with distinct surface charges, including lipid nanoparticles formulated with the commercially available ionizable lipid SM-102 used in COVID-19 vaccine formulations, is applied to ileoids to assess changes in cellular uptake following degradation of surface-associated mucins. Together, these findings establish apical-out ileoids as a physiologically relevant human platform for investigating the ileal epithelial interface and its interactions with microorganisms and nanomaterials.

## 2. Methods

### 2.1. Ileal biopsy collection and ethics collection

Biopsy samples from human ileum were obtained during routine colonoscopies at the endoscopic unit of Uppsala University Hospital, Uppsala, Sweden. Prior to colonoscopy, the project and procedures were described to each participant. If willing, the patient signed an informed consent form, which was retained by the responsible clinicians. In connection with the regular colonoscopy, no more than 4 biopsy samples were taken from the distal small intestine as part of the study, each sample measuring approximately 3-4 mm in diameter. The samples were pseudonymized, placed in buffer for transportation, and then used in the laboratory for ileoid establishment. No identifying patient data (name, personal number, medical history, etc) was used in the research. When samples were no longer required, they were destroyed. The procedures for sample acquisition and handling, as well as generation and experimentation on ileoid cultures, were all approved by the appropriate legal body, Etikprövningsmyndigheten, Sweden: permit number d.nr. 2023-03033-01. The tissue samples were either snap frozen in liquid nitrogen and stored at -150°C, pending global proteomics analysis, or used immediately for isolation of adult ileal stem cell-containing crypts for ileoid establishment.

### 2.2. Human ileoid establishment and maintenance

Resected intestinal biopsy samples were washed in ice-cold phosphate-buffered saline (PBS; Thermo Fisher Scientific/Gibco, Waltham, MA; USA) and cut into micrometer-sized pieces. Epithelial crypts were dissociated after washing in ice-cold PBS using gentle cell dissociation reagent (GCDR) (STEMCELL Technologies, Vancouver, Canada) and nutation at 4°C, followed by trituration. Epithelial fragments were filtered through a 70 µm cell strainer (Corning, Corning, New York, USA) and crypt-enriched fractions resuspended in 50 µL Matrigel (Corning) domes in 24-well plates (Sarstedt, Nümbrecht, Germany). The embedded crypts were cultured in IntestiCult human organoid growth medium (OGM, STEMCELL Technologies, Vancouver, Canada) supplemented with 100U mL^−1^ penicillin-streptomycin (Thermo Fisher Scientific/Gibco) and 10 μM Rho-kinase inhibitor (Rock-i; Y-27632, STEMCELL Technologies) at 37°C and 5% CO_2_. OGM medium was refreshed after 2 days without the Rock-i. Newly established ileoids were cryopreserved at passage 3-4 by gently dissolving Matrigel domes in ice-cold Dulbecco’s modified Eagle’s medium (DMEM)/F12 (Thermo Fisher Scientific/Gibco) with 0.25% bovine serum albumin (BSA, Thermo Fisher Scientific/Gibco) and resuspending the ileal ileoids in DMEM/F12/10% heat-inactivated fetal bovine serum (FBS, Thermo Fisher Scientific/Gibco)/10% DMSO (Sigma-Aldrich, Darmstadt, Germany), followed by freezing in a Mr. Frosty^TM^ container at -80°C overnight and transfer to liquid nitrogen gas phase for long-term cryopreservation.

For maintenance of recently established and thawed ileoids, the cultures were passaged weekly, as previously described in Tzioufa, F. et al, at a ratio of ∼1:5.^[9]^ Briefly, the Matrigel domes were manually broken up by pipetting 20 times with 500 μL GCDR, using 1 mL pipette tips. Materials from 3-4 wells were pooled together, then nutated on a nutating table (20 rpm, FisherBrand, Pittsburg, PA, USA) for 10 minutes, before being centrifuged at 300 × *g* for 5 minutes at 4°C. Pelleted cell fragments were resuspended and centrifuged 1-3 times with 3 mL DMEM/F12/0.25% BSA. Ileoid fragments without Matrigel residuals were again resuspended in 50 µL domes of Matrigel:DMEM/F12/0.25% BSA at a ratio of 4:1, divided over 3 domes per well in 24-well plates, and cultured at 37°C and 5% CO_2_. OGM medium containing 100 U mL^−1^ penicillin-streptomycin was refreshed every 2-4 days. Ileoids established from four different donors (pseudonym as follows, donor 1: Hu23-01il, donor 2: Hu23-02il, donor 3: Hu23-03il, donor 4: Hu24-01il) were used. Ileal biopsies from donor 4 served as the reference native biopsies for proteomic analysis.

### 2.3. Ileoid suspension culture and harvesting for experiments

Culture medium at day 3 post-splitting was removed and exchanged with ice-cold Cell Recovery Solution (Corning). Matrigel domes were gently lifted up using 1 mL pipette tip and transferred within the solution to another 15 mL Falcon tube. Material from 3-4 wells was pooled together and incubated on ice for 1 hour on a nutating table (20 rpm, FisherBrand). Supernatant was carefully aspirated, and ileoid pellets were washed with 3 mL of ice-cold DMEM/F12/0.25% BSA. Following centrifugation (300 x *g*, 5 minutes at 4°C), supernatants were removed, and the pellets were resuspended in IntestiCult human organoid differentiation medium (ODM; STEMCELL Technologies), supplemented with DAPT (STEMCELL Technologies) and 100 U mL^−1^ penicillin-streptomycin. Ileoid suspensions (500 μL per well) were transferred into pre-cooled 24-well ultra-low attachment plates (Corning). To maintain the basal-out polarity of ileoids in suspension culture, Matrigel was added to the medium at a final concentration of 7.5%. For apical-out polarity reversal, Matrigel was omitted as previously described.^[10,40]^ All suspension cultures were incubated at 37°C in a humidified atmosphere containing 5% CO_2_, with medium replacement every other day. Before experiments, apical-out and basal-out ileoids were strained and washed with PBS in 25 µm or 40 µm mini cell strainers (Funakoshi, Tokyo, Japan), respectively, then resuspended in the desired solution.

### 2.4. Immunocytochemistry

Apical-out and basal-out ileoids were collected in PBS in Protein LoBind tubes (Eppendorf, Hamburg, Germany) pre-treated with PBS containing 0.1% BSA. The ileoids were then fixed for 30 minutes at room temperature with a paraformaldehyde solution (PFA) at pH 7.4, consisting of 2% PFA (Thermo Fisher Scientific) in PBS. Following fixation, ileoids were washed three times with PBS for 10 minutes each. Ileoids were then permeabilized with 0.1% saponin (Thermo Fisher Scientific) and blocked with 10% goat serum (Abcam Limited, Cambridge, UK) in PBS, each for 30 minutes at room temperature.

Primary antibodies used included anti-zonula occludens-1 (ZO-1, 1:100, mouse monoclonal, Thermo Fisher Scientific), anti-SLC27A4/FATP4 (1:100, rabbit monoclonal, Thermo Fisher Scientific). These antibodies were prepared in a blocking/permeabilization buffer containing 1% goat serum, 1% saponin, and 1% Triton X-100 (Sigma-Aldrich, Darmstadt, Germany) at the desired concentrations. Ileoids were resuspended in 150 µL of the primary antibody staining solution and incubated overnight at 4°C. Following incubation, ileoids were washed three times with PBS for 10 minutes each.

To detect the proteins of interest, fluorescence-labelled secondary antibodies were used: Alexa Fluor 488 (1:500, anti-mouse, Abcam), Alexa Fluor 488 (1:500, anti-rabbit, Thermo Fisher Scientific). Phalloidin conjugated with Alexa Fluor 660 (1:400, Thermo Fisher Scientific), and wheat germ agglutinin (WGA) conjugated with Alexa Fluor 488 (1:50, Thermo Fisher Scientific) were also added to stain actin and GlcNAc residues, respectively. Ileoids were resuspended in 150 µL of secondary antibody and phalloidin solution, with or without WGA depending on the experimental design, and incubated overnight at 4°C. After three washes with PBS, each for 10 minutes, the ileoids were stained with DAPI (Sigma-Aldrich) in blocking/permeabilization buffer for 20 minutes. Ileoids were then washed 2 times with PBS, 10 minutes each, subsequently resuspended in 20 µL of PBS and transferred to a microscopy glass slide (KEBO-Lab, Stockholm, Sweden). Excess PBS was removed, and DAKO fluorescent mounting medium (Agilent, Santa Clara, CA, USA) was added to the sample. A coverslip, with vacuum grease applied at the corners as spacers, was placed on top of the slide. The ileoids were imaged using a Leica Stellaris 5 confocal microscope (Leica, Wetzlar, Germany) equipped with a 10x/0.4 air objective, a 40x/0.95 air objective and a 63x/1.4 immersion oil objective.

### 2.5. Quantitative global proteomic analysis of ileoids and ileal biopsies

Apical-out and basal-out ileoids were washed twice with ice-cold PBS and pooled (3 wells per sample) into Protein LoBind tubes (Eppendorf). Following centrifugation, supernatants were carefully removed and samples were washed twice more with ice-cold PBS. Ileoids were then pelleted and snap-frozen in liquid nitrogen, before being stored at -80°C freezer. On the day of preparation, samples were prepared as described previously.^[41]^ In brief, frozen samples were rapidly thawed, lysed, and reduced in 25 mM HEPES buffer (pH 7.6, Thermo Fisher Scientific) containing 4% sodium dodecyl sulfate (Sigma-Aldrich) and 1 mM dithiothreitol (Sigma-Aldrich). Frozen biopsies were rapidly thawed, washed with ice-cold PBS, manually homogenized, and subsequently reduced in the same lysis buffer. Protein extraction and digestion were carried out using a modified single-pot, solid-phase-enhanced sample-preparation (SP3) protocol^[42]^ with overnight digestion by trypsin (1:100, Thermo Fisher Scientific) and lysyl endopeptidase (Lys-C, 1:100, Thermo Fisher Scientific).

The following day, peptide digests were purified on magnetic beads (Cytiva, MA, USA) and subsequently eluted via a hydrophobicity shift, transferred into glass vials, and dried in a SpeedVac (Thermo Fisher Scientific) at 40°C. Peptides were separated on an EASY-Spray C18 column (50 cm × 75 µm inner diameter, Thermo Fisher Scientific) with a 300 nL minute^−1^ acetonitrile/water gradient (0.1% formic acid) and analyzed on an Orbitrap Q Exactive HF mass spectrometer (Thermo Fisher Scientific). Raw MS data were processed in MaxQuant v2.6.2.0^[43]^ against the UniProtKB/Swiss-Prot human and mouse reference database curated on 2024-12-03^[44]^.

Data analysis was performed using an in-house proteomics pipeline in R (version 4.2.3). Differential expression analysis was conducted with DEqMS^[45]^ (adjusted p-value ≤ 0.05; absolute log_2_ fold-change ≥ 1), then inputted into gene set enrichment analysis (GSEA; p-value ≤ 0.05; Benjamini-Hochberg correction) performed with clusterProfiler^[46]^ (version 3.18) in R. Protein abundances (fmol µg^−1^ total protein) were estimated using the Total Protein Approach^[47]^. For high-confidence quantification, proteins were considered quantified only when identified by a minimum of three unique and razor peptides. Mass spectrometry proteomics data were deposited to the ProteomeXchange Consortium through the PRIDE partner repository^[48]^ and are available under the dataset identifier PXD084272.

### 2.6. Brightfield microscopy

Apical-out and basal-out ileoids were collected in PBS in an ultra-low attachment plate. Brightfield imaging was performed at 4x/0.1, 10x/0.25 and 20x/0.4 magnifications using a Nikon Eclipse TS100 Inverted Routine Microscope with a Nikon DS-Fi2 camera mount (Nikon Corporation, Tokyo, Japan). Morphological properties of the organoid’s middle-cross-section, including area, Feret’s diameter and circularity, for both apical-out and basal-out ileoids, were approximately quantified using Fiji (version 1.54p)^[49]^ on brightfield images at 4x magnification.

### 2.7. Scanning electron microscopy

Scanning electron microscopy was performed with minor modifications to a previously reported procedure.^[9]^ In brief, ileoids were fixed with 2.5% glutaraldehyde in 0.1 M PHEM-buffer [60 mM piperazine-N,N’-bis (2-ethanesulfonic acid) (Sigma-Aldrich), 25 mM HEPES (4-(2-hydroxyethyl)-1-piperazineethanesulfonic acid (Sigma-Aldrich), 10 mM egtazic acid (Sigma-Aldrich), and 4 mM MgCl_2_ x 6H_2_0 (Sigma-Aldrich), pH 6.9] overnight at 4°C. Samples were allowed to adhere to poly-D-lysine coated cover slips (Ted Pella, Redding, CA, USA) overnight. Cover slips were washed and dehydrated with a graded series of ethanol (Sigma-Aldrich) using a Biowave pro+ microwave (Pelco, Ted Pella) and subsequently critically point dried (Leica EM300). Cover slips were then mounted with carbon tape on an aluminium stub and coated with 5 nm of platinum (Quorum Q150T ES, Quorum Technologies, Lewes, UK). The sample morphology was analyzed by field-emission scanning electron microscopy (FESEM; Carl Zeiss Merlin) using the Inlens secondary electron detector at an accelerating voltage of 5 kV and probe current of 120 pA.

### 2.8. Transmission electron microscopy

For general ultrastructural characterization, the method used was adapted from Tzioufa F et al.^[9]^ with minor modifications. In brief, ileoids were fixed with 2.5% glutaraldehyde in 0.1 M PHEM-buffer overnight at 4°C. Samples were washed in buffer and post-stained in 0.8% potassium ferricyanide (Sigma-Aldrich), 1% OsO4 (TAAB Laboratories, Berkshire, UK) in buffer, further stained with 1% tannic acid (Electron Microscopy Sciences, Hatfield, PA) in Milli-Q water and 1% uranyl acetate (TAAB Laboratories), and further dehydrated using a graded series of ethanol. Samples were infiltrated using a Hard grade embedding resin (TAAB Laboratories) and finally polymerized overnight in an oven at 65°C. All sample preparation except fixation was done using a Biowave Pro+ processing microwave (Pelco, Redding, CA). Ultrathin sections of 70 nm were picked up on formvar coated grids and analyzed using a Talos L120C (FEI Company, Eindhoven, The Netherlands) operating at 120 kV. Micrographs were acquired with a Ceta 16M CCD camera (FEI Company) using Velox software (version 2.14.1.40, Thermo Fisher Scientific). Quantification of glycocalyx presence was performed using Fiji (version 1.54j)^[49]^. Glycocalyx contrast intensity was determined by measuring mean gray value of the glycocalyx area on top of each microvillus after normalization to the mean gray value of the local background of the images (n = 63-67).

For the visualisation of endocytic pathways, sample preparation was performed using modifications of previous methods.^[50]^ Briefly, ileoids were initially fixed in 2.5% glutaraldehyde in PBS overnight at 4°C and then washed with PBS before storage at 4°C. They were then treated sequentially with a mixture of potassium ferricyanide and osmium tetroxide in ddH2O, thiocarbohydrazide in ddH2O, osmium tetroxide in ddH2O, lead aspartate (pH 5.5), and uranyl acetate in ddH2O. Each step was irradiated in a microwave (Pelco) at 80W for 3 min, with thorough ddH2O washes in between. Ileoids were then serially dehydrated with ethanol before resin embedding.

For HRP uptake experiments, ileoids were incubated in ODM containing DAPT, 100 U mL^−1^ penicillin-streptomycin, and 10 mg mL^−1^ HRP (Sigma-Aldrich) for 20 min at 37°C and 5% CO₂. Ileoids were then washed with ice-cold PBS, fixed overnight in 2.5% glutaraldehyde in PBS at 4°C, washed with PBS, and stored at 4°C until further processing. Ileoids were then further processed for DAB detection and Epon embedding as described previously.^[51]^ Sections were acquired using a UC6 ultramicrotome (Leica) and ultrathin sections viewed using a JEM-1400 transmission electron microscope operating at 80kV equipped with a Phurona (EMSIS, Münster, Germany) camera.

### 2.9. Lucifer yellow barrier integrity assays in live ileoids

Barrier integrity of live ileoids was assessed using Lucifer yellow (LY) dilithium salt (Biotium; Ex/Em 428/536 nm).^[9]^ Following three washes in DMEM/F12, ileoids were resuspended in DMEM/F12 supplemented with 100 μM LY and incubated for 60 min at 37°C to allow dye loading. The ileoids remained in the LY-containing medium for the duration of the assay. Live-cell time-lapse images were acquired at 60, 90 and 120 minutes following dye addition, with samples maintained at 37°C and 5% CO_2_ throughout imaging. To generate a permeabilized control, EDTA was added to a final concentration of 2 mM at 60 minutes (t = 0) after dye loading and live-cell time-lapse images were acquired in 30-minute interval. Images were acquired on a Nikon Ti2 microscope equipped with a 10X/0.45 NA Plan Apo Lambda air objective (pixel size 1.12 µm) and a back-lit sCMOS camera (Prime 95B, Photometrics). Fluorescence was excited using the blue channel on a Spectra-X light engine (438/29 nm, Lumencor). A quadruple bandpass dichroic and emission filter (Chroma 89402bs & 89402m) was used for LY detection. LY fluorescence was quantified in Fiji (version 1.54j)^[49]^. The barrier integrity was evaluated by comparing the mean gray value measured across the entire ileoid within a consistent focal plane with the mean gray value of the adjacent local background (mean grey value of ileoid over background; n = 42 ileoids per time point).

### 2.10. Nutrient uptake assays with fatty acid analogue

Ileoids were washed thrice with warm DMEM/F12, resuspended in DMEM/F12 containing 2.5 μM C1-BODIPY-C12 (Thermo Fisher Scientific) and 5 μM BSA, and incubated for 10 minutes at 37°C.^[9]^ Then, the ileoids were washed and resuspended in DMEM/F12 with HCS LipidTOX™ Deep Red Neutral Lipid Stain (Thermo Fisher Scientific). Live-cell imaging was performed using a 60x/0.7 NA air objective in custom-built microscope (37°C, 5% CO_2_) to visualize accumulation of C1-BODIPY-C12 within lipid droplets.

### 2.11. Effect of surface-associated mucin removal by StcE on bacterial invasion

#### 2.11.1. Surface-associated mucin removal by StcE

Apical-out ileoids at day 6 of differentiation were washed and resuspended in warm HBSS without or with mucinase StcE protease at different concentration ranging from 2 to 8 μg mL^−1^ (Sigma-Aldrich) in Protein LoBind tubes (Eppendorf), and incubated for 2 hours at 37°C with 5% CO_2_. The ileoids were subsequently passed through mini cell strainers, and the resulting cell pellets were characterized to select the optimal working concentration of mucinase StcE enzyme.

#### 2.11.2. Viability of ileoids following treatment with StcE

Apical-out ileoids with and without StcE treatment were mixed subsequently with the lysis buffer Reagent A100 (ChemoMetec, Allerod, Denmark), then stabilizing buffer Reagent B (ChemoMetec). Processed samples were loaded into NucleoCassettes (ChemoMetec) and analyzed on the Nucleocounter NC-100 (ChemoMetec), which measured both total and non-viable cell concentrations to determine organoid viability.

#### 2.11.3. Live imaging of ileoids in presence of StcE

Apical-out ileoids with and without StcE treatment were stained with the solution containing Ulex europaeus agglutinin 1 (UEA-1) conjugated with fluorescein dye (UEA-1-FITC, Vector Laboratories, Newark, CA, USA) for 30 minutes at 37°C with 5% CO_2_. Following careful removal of the supernatants, the ileoids were washed twice with HBSS, and subsequently transferred to µ-Slide I Luer 0.4 mm chambers (ibidi GmbH, Gräfelfing, Germany) for live-cell imaging. The fluorescence intensity of UEA-1-FITC (Ex/Em 495/514 nm) was utilized as a marker to evaluate the retention of mature surface-attached mucin residue in the ileoid glycocalyx, thereby mucin removal activity of the StcE enzyme within the ileoids. StcE concentration with the most promising results relating to mucin degradation capability and ileoid viability was chosen for further experiments.

#### 2.11.4. Bacterial invasion assays

Apical-out enteroids with and without StcE treatment at optimal concentration (8 ug mL^−1^) were mixed in ODM supplemented with caspase inhibitor benzyloxycarbonyl-valine-alanine-aspartate-fluoromethylketone (Z-VAD-FMK, AH Diagnostics, Tilst, Denmark), then transferred into 24-well ultra-low attachment plates. For bacteria preparation, the *Salmonella enterica* serovar *Typhimurium* strain SL1344 carrying a vacuolar reporter plasmid p*ssaG*-GFP (*S*.Tm *wt/*p*ssaG-*GFP reporter strain)^[52–54]^ was grown for 12 hours in LB/0.3 M NaCl containing appropriate antibiotics, diluted 1:20, and subcultured for 4 hours in 3 ml of the same medium without antibiotics. Inocula were cultured in a roller drum incubator. Strains were diluted in ODM + 100µM Z-VAD-FMK to achieve the desired MOI.

For infection, ileoids were exposed to *S*.Tm *wt/*p*ssaG-*GFP reporter strain by adding a diluted inoculum to each well at a multiplicity of infection (MOI) of 10. Following a 20-minute incubation at 37°C with 5% CO_2_, ileoids were washed with DMEM/F-12 containing 0.25% BSA, then resuspended in ODM containing gentamicin (Sigma-Aldrich) and Z-VAD-FMK before being transferred to µ-Slide I Luer 0.4 mm (ibidi GmbH) for live-imaging. Images were acquired 2 hours post-infection on a Nikon Ti2 microscope equipped with a 10X/0.45 NA Plan Apo Lambda air objective (pixel size 1.12 µm) and a back-lit sCMOS camera (Prime 95B, Photometrics). Fluorescence was excited using the cyan channel on a Spectra-X light engine (475/34 nm, Lumencor). A quadruple bandpass dichroic (Chroma 89402bs) and a ET525/50m emission filter (Chroma) was used for GFP detection.

To evaluate the invasion efficiency of the bacteria within the ileoids, the number of intracellular *ssaG*-GFP^+^ puncta was measured in each imaged organoid using adaped in-house plugin in Fiji (version 1.54j)^[49]^. Ileoids were manually outlined cross-sectionally at the middle plane as region of interest (ROI). For each manually defined ROI, a 10-µm expanded ROI was generated and the annular region between the original and expanded ROI was used to estimate local background intensity for subsequent normalization of fluorescent intensity of whole ileoids. Images were then contrast-normalized (display range 0-160), thresholded using the Moments algorithm, converted to binary masks, and despeckled to remove isolated noise pixels. Intracellular *ssaG*-GFP^+^ puncta were identified using the Analyze Particles function with a minimum particle size of 15 µm² and no upper size limit.

### 2.12. Effect of surface-associated mucin removal by StcE on nanoparticle transportation

#### 2.12.1. Preparation of lipid nanoparticles

Thin lipid films consisting of DSPC (10 mol%) (850365P, Avanti Research, Sigma-Aldrich), cholesterol (38.5 mol%) (700000P, Avanti Research), SM-102 (36-0102, Larodan AB, Solna, Sweden) (50 mol%), and DMG-PEG2000 (1.5 mol%) (36-6014, Larodan AB) were prepared by preparing individual lipid stocks in chloroform (99+%, Thermo Fisher Scientific). The stocks were then mixed using appropriate volumes dispensed using glass Hamilton syringes to reflect the stated molar ratios. The mixing of the lipid components in chloroform was performed in a borosilicate vial, and the solvent was allowed to evaporate overnight in a fume hood. Vials were then sealed under a stream of nitrogen and stored at -20°C until further use. The lipid nanoparticles (LNPs) were fluorescently labelled by dissolving Cy5-DOPE (810335C, Avanti Research, Sigma-Aldrich) in chloroform at 50 µg mL^−1^ and adding a volume corresponding to 0.1 mol% of the total lipid content to the film. The solvent was evaporated overnight, and vials were sealed under nitrogen and stored at -20°C. The final lipid mass of the lipid films was 2.8 mg.

To formulate LNPs, the lipid film was dissolved in ethanol (99+%, Thermo Fisher Scientific) to a final total lipid concentration of 12.5 mM and loaded into a 2.5 mL gastight glass syringe (Hamilton 1002 TLL). A second glass syringe was loaded with an equal volume of 50 mM sodium citrate buffer (pH 4.0) (buffer prepared using anhydrous tri-sodium citrate and citric acid monohydrate, Merck). The syringes were mounted on a dual-drive syringe pump (Pump 33 DDS, Harvard Apparatus, Holliston, MA, USA) and connected to a passive polycarbonate herringbone mixing chip (CAT:10000019, microfluidic ChipShop GmbH, Jena, Germany). Mixing was performed at a total flow rate of 7 mL minute^−1^ (5.25 mL minute^−1^ for the buffer syringe and 1.75 mL minute^−1^ for the lipid syringe). After activating the pump, the initial 133 μL collected from the syringe pump was discarded and the rest of the sample was collected. After collection from the microfluidic pump, the sample was loaded into Slide-A-Lyzer™ MINI Dialysis Devices, 3.5K MWCO (Thermo Fisher Scientific) (100 µL of LNPs per dialysis cup) and dialyzed overnight against 1000 volumes of PBS (-Ca^2+^, -Mg^2+^) (Gibco) at room temperature, after which the pH was checked using pH strips to confirm that the sample had reached neutral pH 7.4 (Macherey Nagel X100 pH test strip, Thermo Fisher Scientific).

#### 2.12.2. Physicochemical characterisation by nanoparticle tracking analysis and dynamic light scattering

Particle diameter, concentration, zeta potential and population analysis were performed using Nanoparticle Tracking Analysis (NTA) (ZetaView® x30, Particle Metrix GmbH, Germany). Polystyrene and LNPs were measured by performing a 1:250000 and 1:20000 (v/v) dilution in 10% PBS (-Ca^2+^, -Mg^2+^) (Gibco), respectively, and injecting the sample into the instrument using a 1 mL sterile Luer plastipak syringe (Becton Dickinson S.A., Franklin Lakes, NJ, USA). Measurements (concentration, size and zeta potential) were performed sequentially in triplicates using the SC488 laser (no filter) and instrument settings of gain = 40 and shutter = 200. Measurements were performed at 25°C, ensuring that the number of traced particles exceeds 500 (N = 1, n = 3). Presented results from the NTA are reported as median derived from the number-weighted distribution. The hydrodynamic diameter of LNPs was also measured using a Zetasizer Pro (Malvern Panalytical, Malvern, United Kingdom). For size characterization, lipid nanoparticles were diluted 1:200 (v/v) in 1× PBS (-Ca^2+^/-Mg^2+^, Gibco) inside polystyrene microcuvettes (Brand GmbH, Wertheim, Germany). Samples were then equilibrated at 2°C for 120 second inside the machine. Dynamic light scattering (DLS) measurements were performed in triplicate using automatic attenuator selection. To prevent fluorescence interference from labeled LNPs without compromising detection sensitivity, an integrated fluorescence filter was utilized.

#### 2.12.3. Determination of nanoparticle uptake using plate reader

Apical-out ileoids with and without StcE treatment at optimal concentration were resuspended in culture medium containing NPs. The NP used was either 100 nm polystyrene NPs, negatively charged (Thermo Fisher Scientific) or positively charged (Sigma-Aldrich), or nearly neutrally charged LNPs. Polystyrene nanoparticles or LNPs were applied to the apical-out ileoids at the final concentration of 8 × 10^10^ and 2 × 10^10^ particles mL^−1^ in pre-warmed DMEM supplemented with 10% FBS (Thermo Fisher Scientific/Gibco) and 100 U mL^−1^ penicillin-streptomycin (Thermo Fisher Scientific/Gibco), respectively. After 4-hour incubation at 37°C with 5% CO_2_, ileoids were strained, washed three times with PBS, and transferred to Protein LoBind tubes (Eppendorf). They were then resuspended in pre-warmed TrypLE Express (Thermo Fisher Scientific), mixed thoroughly, and incubated for 30 minutes in the incubator to achieve dissociation. Single cells were pelleted, washed three times, resuspended in FACS buffer (PBS + 0.1% BSA, Thermo Fisher Scientific), and analyzed with Tecan Spark plate reader. The excitation and emission wavelengths for the NPs were set to 580 nm and 605 nm, respectively, each with a 10 nm bandwidth filter. Fluorescence intensity of fluorescently labelled NPs at 25°C was used as the marker to evaluate the uptake efficiency of the NPs. Nanoparticle internalization data were analyzed using one-way ANOVA test (n = 3).

### 2.14. Statistical analysis

Statistical analyses were performed using GraphPad Prism 10.0.0 (GraphPad Software, San Diego, CA, USA), unless otherwise stated. For microscopy-based experiments, the sample size (n) corresponded to the number of ileoids analyzed per condition unless indicated otherwise in the figure legends. Data distribution was assessed using appropriate normality tests prior to statistical comparison. Depending on data normality, parametric or nonparametric statistical tests were applied as specified in the respective figure legends. Statistical significance was defined as p < 0.05 and is indicated as follows: *\*p <* 0.05*, **p <* 0.01*, ***p <* 0.001*, and ****p <* 0.0001.

## 3. Results

### 3.1. Apical-out human ileoids recapitulate key characteristics of native ileal tissue

We established a protocol to generate suspension-grown ileoids from healthy human ileal biopsies by adapting the method previously described.^[9,40]^ (Figure 1A). Ileoids cultured in ODM lacking Matrigel adopted an apical-out configuration, whereas those maintained in medium containing low concentrations of Matrigel retained a basal-out configuration. To assess epithelial polarity, ileoids cultured under both conditions were immunostained for F-actin and the tight junction protein ZO-1. Co-localization of F-actin and ZO-1 was used as an indicator of apical membrane localization (Figure 1B).^[10]^ Image analysis revealed that complete removal of Matrigel from the culture medium induced a complete reversal of epithelial polarity, resulting in ileoids with an apical-out configuration (Figure 1C). In contrast, ileoids maintained in Matrigel-containing medium predominantly exhibited a basal-out configuration (96.8%, Figure 1C). Together, these findings demonstrate that our protocol enables robust and reproducible generation of apical-out human ileoids in suspension culture.

**Figure 1.**
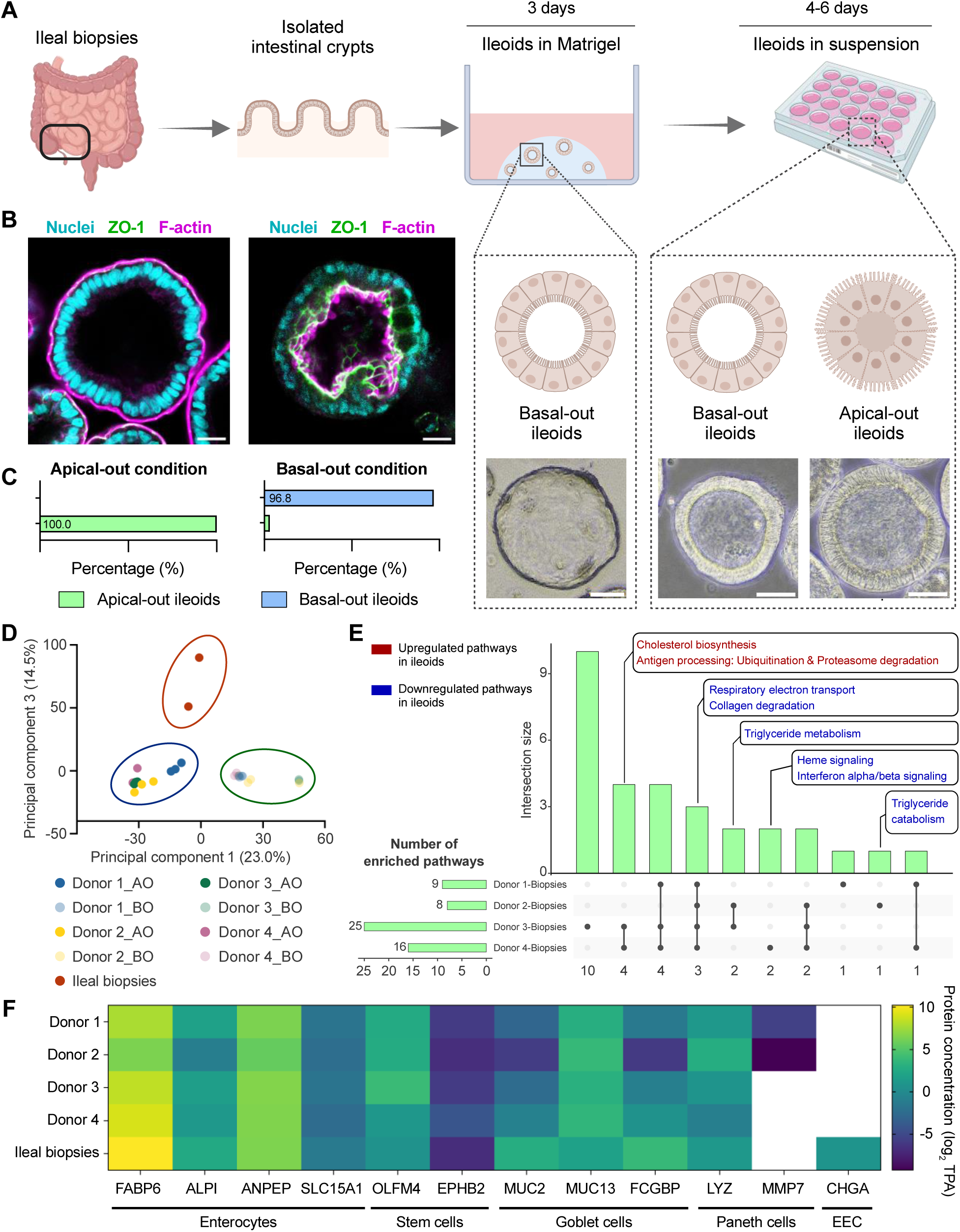
Global proteomic profiling of suspension-grown human ileoids. **(A)** Workflow for establishing human ileoids from intestinal biopsies. Schematic overview of ileoid culture in extracellular matrix (Matrigel) and suspension cultures derived from human ileal biopsies. Representative bright field micrographs of ileoids in Matrigel as well as basal-out and apical-out ileoids in suspension. Scale bars, 100 µm. **(B)** Representative confocal images following staining of zonula occludens-1 (ZO-1, green), phalloidin (F-actin; magenta) and nuclei (DAPI; cyan). Scale bars, 20 µm. **(C)** Quantification of apical-out and basal-out ileoids within suspension cultures by confocal microscopy. Data are shown as frequencies (n = 75-120). **(D)** Principal component analysis of the proteomic profiles of each ileoid line alongside reference ileal biopsy samples. **(E)** Upset plot illustrating the overlap of enriched pathways among ileoid lines compared with reference ileal biopsies. **(F)** Expression patterns of selected cell-type specific marker proteins across ileoid lines and ileal biopsies. Data are shown as log_2_ transformed of protein concentration after from total protein approach (TPA) quantification method.

Subsequently, apical-out ileoids derived from four individual donors were compared to identify the most representative model for subsequent experiments. Morphological and proteomic analyses revealed donor-dependent differences, while epithelial cell-type marker expression was broadly comparable across ileoid lines and consistent with native ileal tissue (Figure S1, Figure 1D–F). Among the four donor lines, ileoids from donor 1 showed the greatest overall proteomic similarity to native ileal biopsies, together with high reproducibility among replicates (Figure S2). Accordingly, donor 1 ileoids are selected for all subsequent experiments.

### 3.2. Apical-out ileoids show overall morphological and proteomic stability during serial passaging

To define a suitable experimental window, ileoid morphology and proteomic stability were assessed during serial passaging. Brightfield microscopy was used to quantify Feret’s diameter and circularity at eight-passage intervals beginning at passage 6 (Figure 2A). Ileoids at passage 14 retained comparable size and shape to those at passage 6. However, by passage 22, a modest but significant reduction in diameter (160.8 ± 73.0 µm in passage 6, compared to 141.1 ± 70.3 µm in passage 22) and increased circularity (0.79 ± 0.18 in passage 6, compared to 0.82 ± 0.17 in passage 22) were observed, suggesting progressive phenotypic alterations with extended passaging (Figure 2B). This morphological shift was corroborated by PCA, which revealed a distinct separation in the proteomic profiles of passage 22 ileoids relative to earlier passages (Figure 2C). Of the 4,224 proteins quantified in the proteomic analysis comparing ileoids at passages 6 and 14, only 235 and 226 proteins were significantly downregulated and upregulated, respectively, at passage 14 (Figure 2D). In comparison, the analysis of ileoids at passages 6 and 22 quantified 3,859 proteins and revealed a markedly greater number of differentially expressed proteins at passage 22, with 451 proteins downregulated and 359 proteins upregulated (Figure 2E). Despite consistent expression levels of most proliferation markers across all timepoints, a significant downregulation of goblet cell differentiation marker (MUC2) was detected in ileoids at passages 14 and 22, suggesting reduced goblet cell representation over time (Figure 2F). Furthermore, stem cell (OLFM4) and enterocyte markers (ALPI) were significantly reduced in passage 22 ileoids cultured at passage 22, in comparison to the ones at passage 6. Despite these changes, apoptosis-related protein markers (CASP3, CASP6, CASP7) remained stable over time, indicating consistent organoid viability across passages (Figure 2E, F). Gene set enrichment analysis comparing passages 14 and 22 with passage 6 identified enrichment of Reactome pathways associated with cell proliferation and senescence exclusively in passage 22 ileoids (Figure 2G). Together, these findings indicate that apical-out ileoids remain broadly stable across serial passages, although some passage-dependent changes become more apparent at later passages. These findings support the use of ileoids across the passages examined, with passages 6–14 providing a relatively stable experimental window.

**Figure 2.**
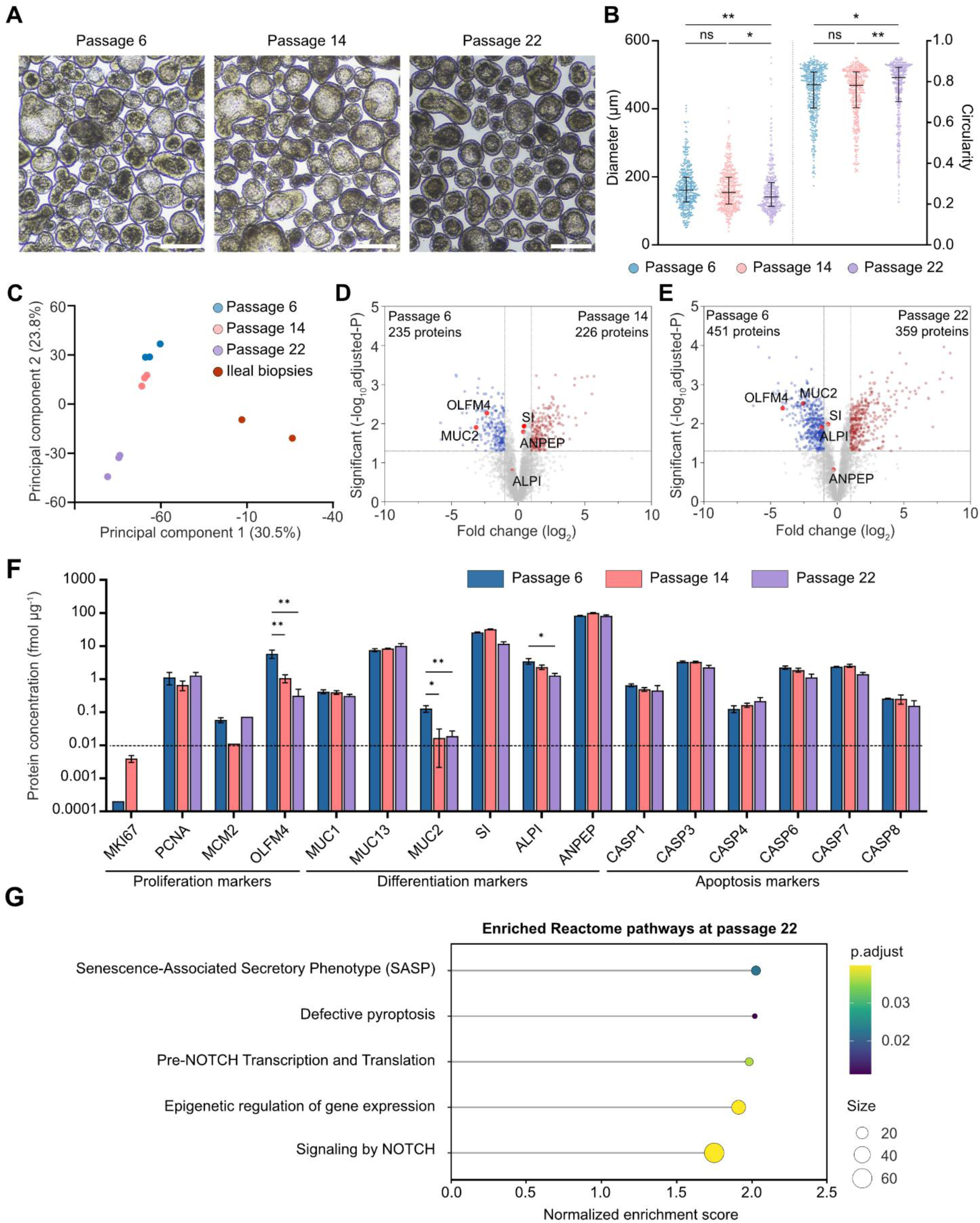
Morphological and proteomic stability of apical-out ileoids during serial passaging. **(A)** Representative brightfield images of ileoids at passages 6, 14, and 22. Scale bars, 200 µm. **(B)** Quantification of Feret’s diameter and circularity of ileoids at passages 6, 14, and 22. Data are shown as median and interquartile range (n = 400). P values were calculated by Kruskal-Wallis test with multiple-comparisons testing; **p < 0.01. **(C)** Principal component analysis of proteomic profiles of ileoids at passages 6, 14, and 22 alongside reference ileal biopsy samples. **(D-E)** Volcano plots showing significantly downregulated and upregulated proteins in ileoids at passage 14 **(D)** and passage 22 **(E)** relative to passage 6. **(F)** Abundance of selected stem cell, differentiation, proliferation, and apoptosis markers across passages 6, 14, and 22. Data are shown as mean ± SD (n = 3). *p < 0.05, **p < 0.01. **(G)** Reactome gene set enrichment analysis comparing passage 22 with passage 6 reveals enrichment of pathways associated with cell proliferation and senescence.

### 3.3. Apical-out ileoids exhibit structural features characteristic of the native ileal epithelium

Scanning and transmission electron microscopy were used to examine the surface morphology and ultrastructure of apical-out ileoids. SEM revealed a dense brush border composed of well-developed microvilli covering the hexagonal apical surface (Figure 3A). TEM confirmed the presence of abundant microvilli, a clearly visible glycocalyx, and tight junctions between adjacent epithelial cells (Figure 3B, S3A). Measurement of average apical surface area (67.66 ± 20.71 µm^2^) and enterocyte height (22.18 ± 4.27 µm, Figure S3B) yielded an estimated enterocyte volume of approximately 1400.00 µm³, consistent with reported *in vivo* values.^[55,56]^ Proteomic profiling further demonstrated broadly comparable expression of key junctional proteins (e.g., TJP1, JAM1, cingulin) in apical-out ileoids and ileal biopsies (Figure 3C). Collectively, these findings indicate that apical-out ileoids closely resemble native ileal epithelium at both the structural and molecular levels.

**Figure 3.**
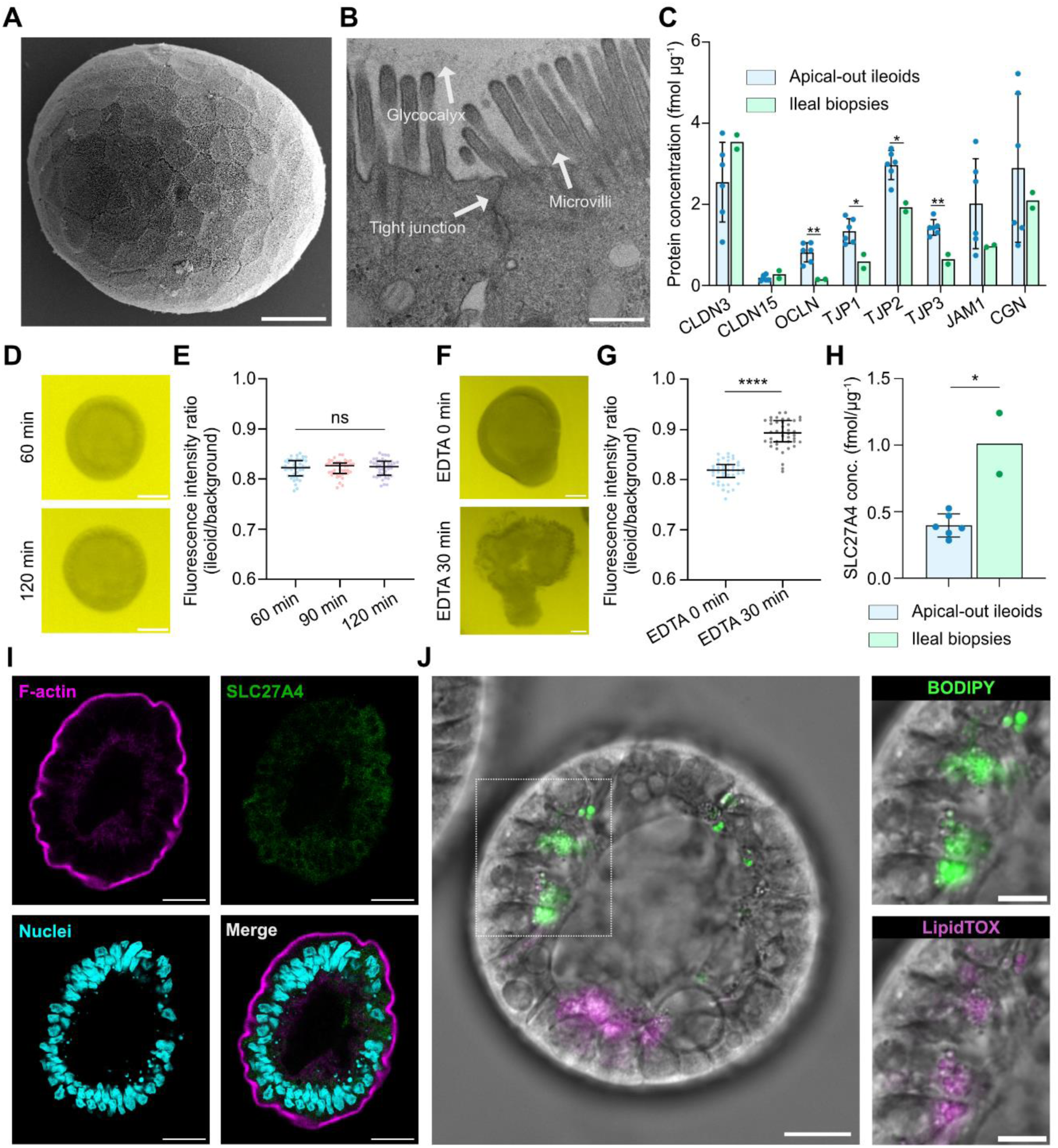
Structural and functional characterization of apical-out ileoids. **(A)** Representative SEM image showing a dense microvillar brush border and hexagonal apical surface architecture. Scale bars, 20 µm. **(B)** Representative TEM micrograph showing well-developed microvilli, glycocalyx, tight junctions. Scale bars, 500 nm. **(C)** Expression of key tight junction proteins in apical-out ileoids as compared to ileal biopsies. Data are shown as mean ± SD where appropriate (n = 6 for ileoids and 2 for ileal biopsies). **(D)** Representative live-cell confocal images of apical-out ileoids incubated with Lucifer Yellow (LY, 100 µM) for 120 minutes demonstrating maintenance of epithelial barrier integrity. Scale bars, 50 µm. **(E)** Quantification of pseudo-luminal LY fluorescence over time. Fluorescence intensity was normalized to background. Data are shown as median with interquartile range (n = 44 ileoids per time point). P values were calculated by Kruskal-Wallis test and no statistical differences were found. **(F)** Representative live-cell confocal images showing increased LY permeability following EDTA treatment. Scale bars, 50 µm. **(G)** Quantification of LY fluorescence intensity within untreated ileoids (incubation with LY for 30 minutes) and ileoids treated with EDTA (additional incubation for 30 minutes with EDTA) was normalized to background. Data are shown as median with interquartile range. An average of 44 observations were analyzed. P values were calculated by Kruskal-Wallis test; ****p < 0.0001. **(H)** Expression levels of SLC27A4 protein (solute carrier family 27 member 4) in apical-out ileoids compared to ileal biopsy in vivo levels. Data are shown as mean ± SD where appropriate (n = 6 for ileoids and 2 for ileal biopsies). **(I)** Representative confocal images following staining with DAPI (nuclei; blue), SLC27A4 (green) and phalloidin (F-actin; magenta) in apical-out ileoids. Scale bars, 20 µm. **(J)** Representative live-cell fluorescence images showing uptake of the fluorescent fatty acid analogue C1-BODIPY-C12 and co-localization with intracellular lipid droplets stained with LipidTOX Deep Red. The dashed box indicates the magnified region. Scale bars, 20 µm; scale bars magnification, 10 µm. Donor 1 ileoids were used throughout.

### 3.4. Apical-out ileoids maintain epithelial barrier integrity and absorptive function

Building upon the observed morphological and proteomic features of the apical-out ileoids, we next assessed their epithelial barrier integrity using the low-permeability tracer lucifer yellow (LY). Co-incubation with LY for 120 minutes did not increase pseudo-luminal fluorescence, indicating that the ileoid barrier remained intact (Figure 3D, E). In contrast, exposure to EDTA for 30 minutes disrupted epithelial barrier integrity, as evidenced by the visible and statistically significant influx of LY from the surrounding medium into the organoid interior (Figure 3F, G). Collectively, these findings demonstrate that apical-out ileoids maintain a physiologically intact barrier under experimental conditions.

The absorptive function of apical-out ileoids was further investigated by examining their ability to take up the fluorescent fatty acid analog C1-BODIPY-C12. Proteomic analysis first confirmed that the fatty acid transporter SLC27A4 was expressed in apical-out ileoids at level within about 2-fold to those observed in ileal biopsy samples (Figure 3H). Immunostaining demonstrated that SLC27A4 was positioned along the apical membrane, supporting its role in facilitating lipid uptake, including C1-BODIPY-C12^[10]^, into the cells (Figure 3I). Following a 30-minute incubation with BODIPY-labeled fatty acid, live cell imaging revealed clear co-localization of the fluorescent probe with LipidTOX™ Deep Red staining intracellular lipid droplets, consistent with cellular uptake of the fluorescent fatty acid analog (Figure 3J). Taken together, these findings show that apical-out ileoids maintain an intact physiological barrier while exhibiting fundamental nutrient absorptive functions characteristic of the small intestine.

### 3.5. Apical-out ileoids retain physiologically relevant endocytic pathways and uptake activity

We next evaluated whether apical-out ileoids provide a suitable model for investigating endocytic uptake from the intestinal lumen by characterizing their endocytic machinery. The expression of key protein markers representing major endocytic pathways was first analyzed in apical-out ileoids and compared with reference ileal biopsies. Proteomic profiling demonstrated expression of core components of the dynamin-dependent endocytic pathways, including fast endophilin-mediated endocytosis (FEME) and clathrin-mediated endocytosis (CME), at levels comparable to those in native ileal biopsies (Figure 4A). Immunostaining revealed that key FEME and CME proteins were localized predominantly to the cortical actin network and plasma membrane of the ileoids (Figure 4B, C). Although markers of caveolae-mediated endocytosis were present in apical-out ileoids, their expression levels were markedly lower than those observed in biopsies (Figure 4A). Additionally, CAV1 localization was identified in lateral and to some extent basal membrane domains (Figure 4D). Proteomic profiling also demonstrated expression of proteins associated with dynamin-independent pathways, including macropinocytosis and CLIC/GEEC-mediated endocytosis (Figure S4A). CTBP1, a central regulator of macropinocytosis, was primarily detected within nuclear and perinuclear regions by immunostaining techniques (Figure S4B). Electron microscopy analysis further revealed membrane pits, narrow subapical tubular structures, and intracellular ring-shaped membrane profiles (Figure 4E, S4C–D). The tubular structures were morphologically consistent with fast endophilin-mediated endocytosis (FEME)^[57]^ or CLIC/GEEC-like intermediates, while the ring-shaped membrane profiles were consistent with CLIC/GEEC-like structures^[58]^. Together with the proteomic identification of components associated with multiple endocytic pathways, these observations provide morphological evidence for diverse endocytic processes in the apical-out ileoid epithelium, including both clathrin-dependent and clathrin-independent mechanisms.

**Figure 4.**
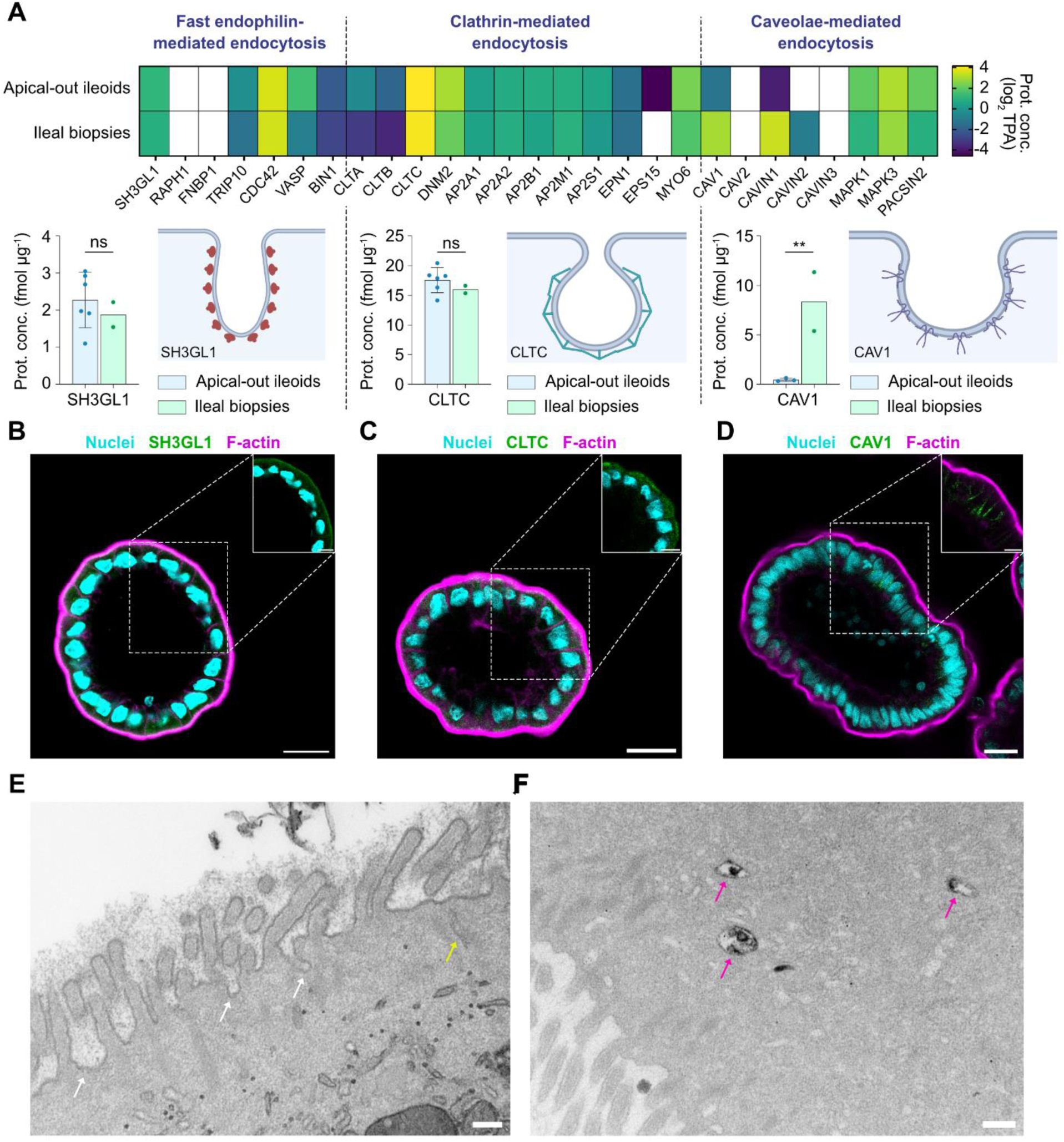
Expression and functional activity of endocytic pathways in apical-out ileoids. **(A)** Expression levels of representative protein markers of clathrin-mediated, fast endophilin-mediated, and caveolae-mediated endocytic pathways in apical-out ileoids compared with reference ileal biopsy samples. Bar graphs are shown as mean ± SD where appropriate (n = 6 for ileoids and 2 for ileal biopsies). **p < 0.01. **(B-D)** Representative confocal images showing localization of proteins involved in **(B)** fast endophilin-mediated endocytosis, **(C)** clathrin-mediated endocytosis, and **(D)** caveolae-mediated endocytosis in apical-out ileoids. Scale bars, 20 µm; scale bars magnification, 10 µm**. (E)** Representative TEM images showing the subapical region containing membrane invaginations/pits (white arrows) and narrow tubular (yellow arrows). These structures are morphologically consistent with endocytic intermediates, including clathrin-associated pits and FEME- or CLIC/GEEC-like tubular structures. Scale bars, 200 nm. **(F)** Representative TEM images demonstrating intracellular uptake of HRP following 20-minute incubation, indicating active fluid- phase endocytosis. Electron-dense diaminobenzidine reaction product is visible within intracellular vesicles (magenta arrows). Scale bars, 200 nm. Donor 1 ileoids were used throughout.

To evaluate the functional activity of the endocytic machineries expressed in apical-out organoids, horseradish peroxidase (HRP), a fluid-phase uptake marker, was applied and analyzed using electron microscopy. Although relatively few clathrin-coated pits were clearly distinguishable, HRP uptake experiments demonstrated abundant intracellular endocytic vesicles (Figure 4F), indicating active endocytic transport despite the seemingly low abundance of morphologically identifiable coated intermediates. Together, these findings establish apical-out ileoids as a physiologically relevant platform for investigating endocytic uptake of luminal cargo in the human distal small intestinal epithelium.

### 3.6. The apical glycocalyx forms a mucin-rich barrier at the ileoid surface

As a next step, we assessed the performance of our model in proof-of-concept experiments using polystyrene nanoparticles (PS NPs) and bacteria. Although the intestinal glycocalyx plays a critical role in restricting the interaction and penetration of such external agents, current understanding of its cellular characteristics remains limited. Because the molecular composition of the intestinal glycocalyx remains incompletely characterized, we first assessed the expression of established glycocalyx components, including representative proteoglycans and membrane-associated mucins.^[28–31]^ Proteomic analysis revealed that crucial proteoglycan syndecan-1 (SDC1) and the CD44 antigen were expressed in apical-out ileoids at levels comparable to those observed in ileal biopsies. In contrast, membrane-associated mucins were expressed at higher levels in the ileoids, with MUC13 showing the most pronounced enrichment (Figure 5A). Glycocalyx localization was confirmed by staining with wheat germ agglutinin (WGA), which binds N-acetylglucosamine (GlcNAc) and sialic acid (Sia) residues present on glycocalyx glycoproteins and proteoglycans.^[59]^ Imaging demonstrated a GlcNAc and Sia-rich layer situated above the F-actin filament layer, thereby confirming the presence and localization of the glycocalyx in the ileoids (Figure 5B).

**Figure 5.**
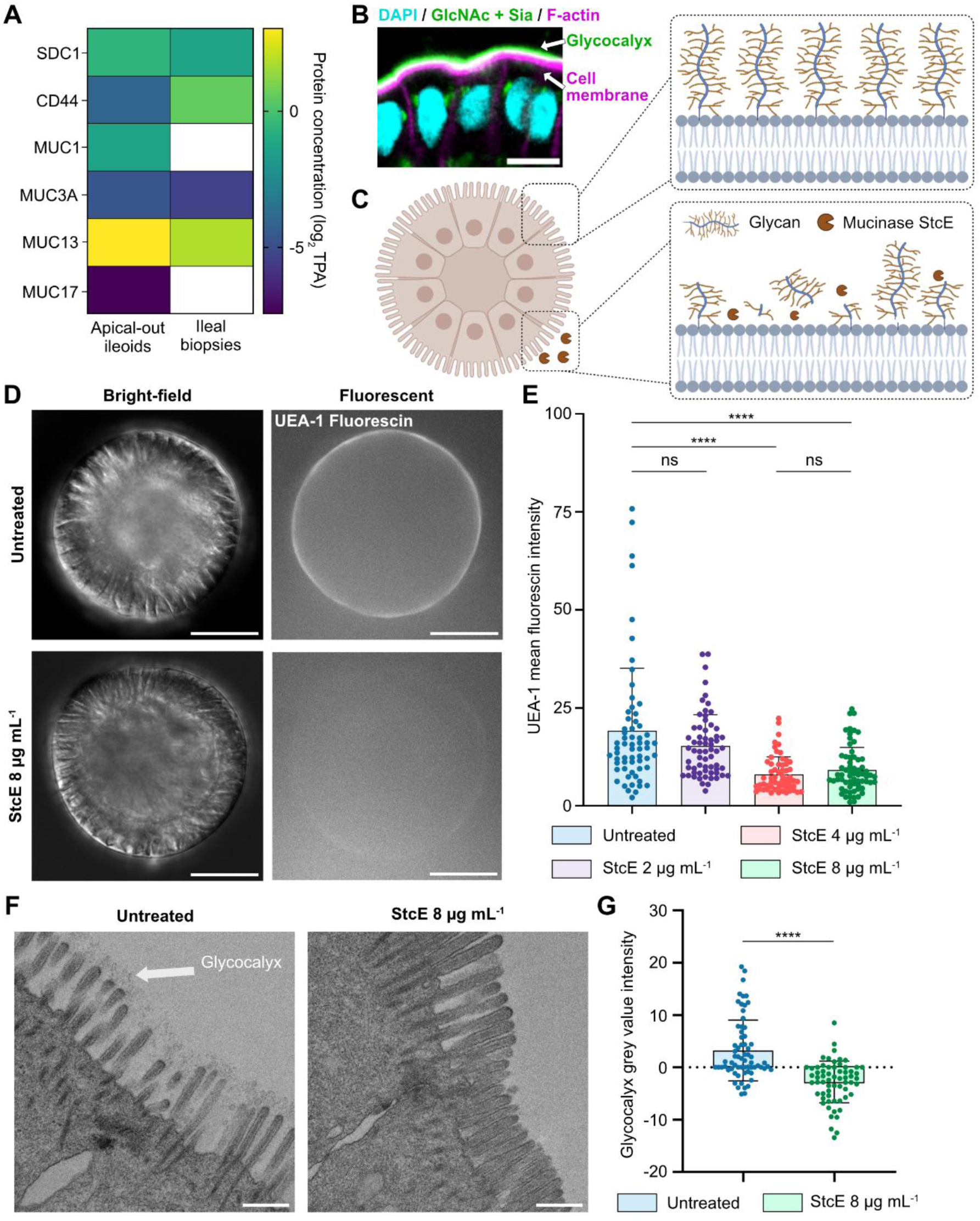
Enzymatic removal of mucin components from the glycocalyx barrier. **(A)** Expression levels of selected proteoglycans and glycoproteins within the glycocalyx of apical-out ileoids compared with reference ileal biopsy samples. **(B)** Representative confocal images showing WGA staining (N-acetylglucosamine and sialic acid residues) overlying the apical F-actin layer, confirming the presence of the glycocalyx. Scale bars, 10 µm. **(C)** Schematic illustration of mucin cleavage by the bacterial metalloprotease StcE. **(D)** Representative live-cell confocal images of UEA-1 staining following treatment with increasing concentrations of StcE. Scale bars, 50 µm. **(E)** Quantification of mean UEA-1 fluorescence intensity after StcE treatment. Data are shown as mean ± SD (n = 60-65). P values were calculated by Kruskal-Wallis test; ****p < 0.0001. **(F)** Representative scanning electron micrographs showing the apical glycocalyx before and after StcE treatment. Scale bars, 500 nm**. (G)** Quantification of glycocalyx electron density (or grey value) following StcE treatment. Data are shown as mean ± SD (n = 63-67). P values were calculated by Mann-Whitney test; **** p < 0.0001. Donor 1 ileoids were used throughout.

We next investigated the role of surface-attached mucin glycoproteins, a major component of the intestinal glycocalyx, in protecting ileoids from external factors. To specifically disrupt the mucin layer, we employed StcE, a bacterial protease derived from *Escherichia coli*, which cleaves mucin domains from the glycocalyx (Figure 5C).^[60,61]^ The optimal StcE concentration was determined using confocal live microscopy with UEA-1 staining, which binds to terminal α-L-fucose residues within mucins. First, none of the tested StcE concentrations had a significant effect on cell viability (Figure S5). Notably, treatment with 4 and 8 µg mL^−1^ StcE markedly reduced UEA-1 staining at the apical membrane of apical-out ileoids compared with untreated controls (Figure 5D, E), indicating efficient removal of mucin components from the apical glycocalyx. SEM further supported this observation, showing reduced mean grey value of the glycocalyx layer in treated ileoids (Figure 5F, G). Based on these findings, 8 µg mL^−1^ StcE is selected as the optimal concentration for mucin removal in subsequent experiments.

### 3.7. Removal of surface-attached mucins promotes bacterial invasion and enhances charge-dependent nanoparticle uptake

To investigate the protective role of mucin components within the glycocalyx barrier of apical-out ileoids, 3D cellular models with intact or enzymatically removed mucins were exposed to external challenges, including invasive bacterial *Salmonella* Typhimurium (*S.*Tm) strain and a panel of nanoparticles with different physicochemical properties. First, to assess the protective function of the glycocalyx barrier in ileoids against bacterial invasion and expansion, we utilized the *S.*Tm wt/p*ssaG*-GFP reporter strain, which expresses GFP upon entry into host cells.^[15,52,54]^ Based on previous reports demonstrating that the glycocalyx limits bacterial invasion^[16,34,62]^, we hypothesized that partial removal of mucin components would facilitate invasion by the *S.* Tm wt*/*p*ssaG*-GFP reporter strain, resulting in increased intracellular GFP expression (Figure 6A). Indeed, StcE-treated ileoids exhibited significantly more *ssaG*-GFP^+^ puncta foci (2.77 ± 3.35 puncta foci in StcE-treated ileoids compared to 1.34 ± 1.98 puncta foci in the control condition), indicating greater augmented invasion by the *S.* Tm wt*/*p*ssaG*-GFP reporter strain in the ileoids with a degraded mucin-linked glycocalyx expression (Figure 6B-C).

**Figure 6.**
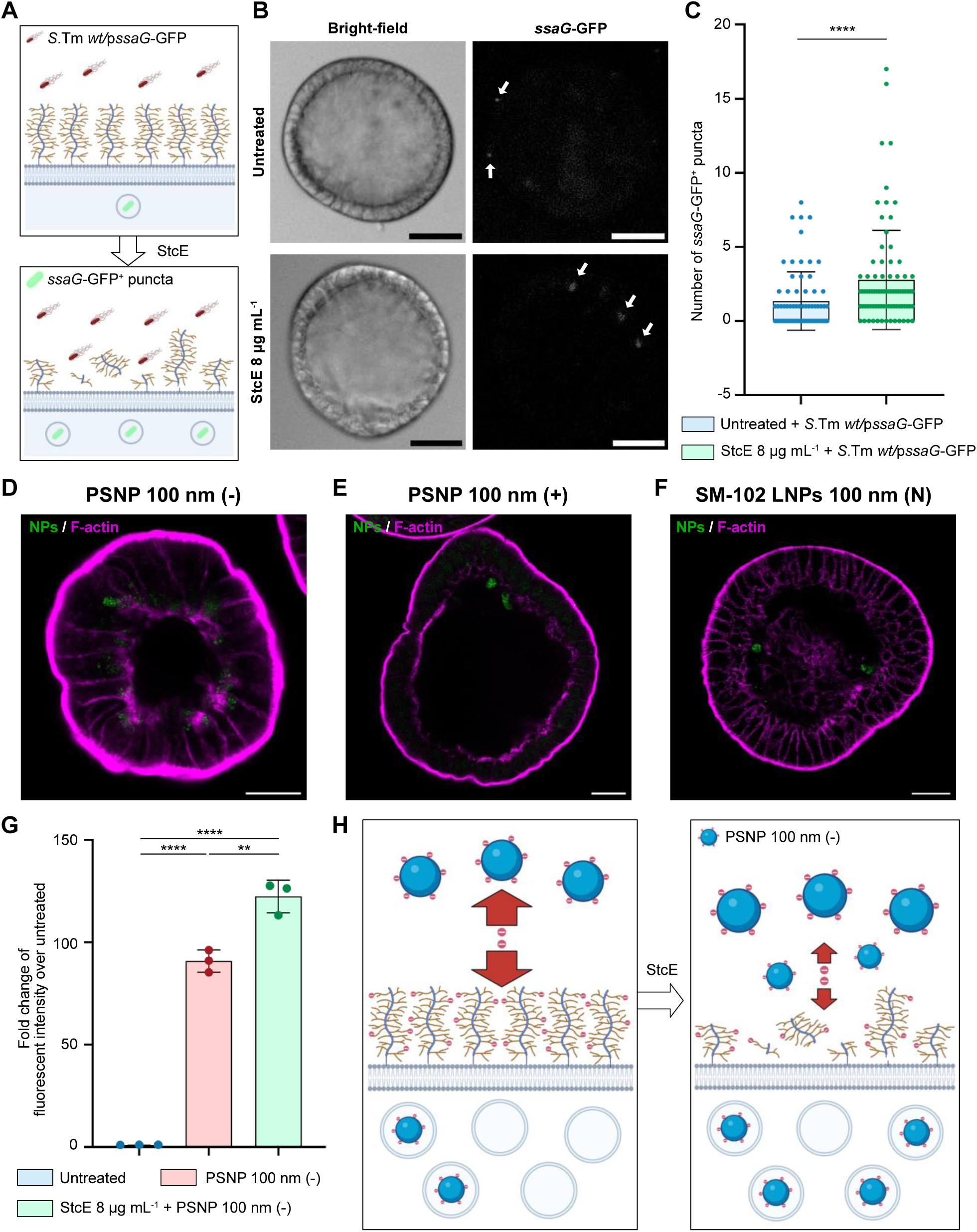
Mucin depletion enhances bacterial invasion and charge-dependent nanoparticle uptake. **(A)** Schematic overview of the experimental design illustrating StcE-mediated mucin removal followed by infection with S. Tm wt/pssaG-GFP. **(B)** Representative live cell images showing increased intracellular ssaG-GFP^+^-bacteria puncta (white arrow) in StcE-treated ileoids compared with untreated controls. Scale bars, 50 µm. **(C)** Quantification of intracellular ssaG-GFP^+^ puncta per ileoid. Data are shown as mean ± SD (n = 72-82). P values were calculated by Mann-Whitney test; ****p < 0.0001. **(D–F)** Representative confocal images showing cellular uptake of **(D)** carboxylate-modified polystyrene nanoparticles, **(E)** amine-modified polystyrene nanoparticles, and **(F)** SM-102 lipid nanoparticles by ileoids with an intact glycocalyx barrier. Scale bars, 20 µm. **(G)** Quantification of intracellular nanoparticle fluorescence intensity in untreated and StcE-treated ileoids. Carboxylate-modified nanoparticles exhibited significantly increased uptake following mucin removal, whereas uptake of amine-modified polystyrene nanoparticles and SM-102 lipid nanoparticles was unchanged. Data are shown as mean ± SD (n = 3). P values were calculated by one-way ANOVA test with multiple-comparisons testing; **p < 0.01, ****p < 0.0001. **(H)** Proposed model illustrating the charge-selective barrier function of the ileal glycocalyx. Partial removal of negatively charged mucin components preferentially enhances uptake of negatively charged nanoparticles. Donor 1 ileoids were used throughout.

Having established that the apical ileoid glycocalyx acts as a protective barrier against bacterial invasion, we next asked whether it also regulates interactions between the intestinal epithelial surface and nanoparticles. To investigate whether these interactions were influenced by nanoparticle surface charge, we exposed mucin-intact and mucin-depleted degraded ileoids to nanoparticles with distinct surface charge properties. Negatively charged (carboxylate-modified) and positively charged (amine-modified) polystyrene nanoparticles with a mean diameter of 100 nm were used. Near-neutral particles were represented by LNPs formulated with a lipid composition which is analogous to those used in the Moderna mRNA COVID-19 vaccines and therefore includes the ionizable lipid SM-102.^[63]^ Nanoparticle tracking analysis confirmed that all nanoparticle types exhibited comparable hydrodynamic diameters (91.1 – 107.7 nm), and their surface charges aligned with manufacturer specifications (Figure S6A). In addition, the Z-average hydrodynamic diameter of the formulated LNPs was 107 ± 1 nm with a polydispersity index (PDI) of 0.12 (Figure S6B). Initial uptake experiments demonstrated that all nanoparticle formulations were internalized by apical out ileoids following a 4-hour incubation, as evidenced by immunofluorescence imaging (Figure 6D-F). Building on these baseline findings, we next evaluated how enzymatic removal of surface-attached mucins influences charge-dependent nanoparticle uptake. We hypothesized that partial loss of negatively charged mucin components would preferentially enhance uptake of anionic nanoparticles by reducing electrostatic repulsion, whereas neutral or cationic particles would not gain a similar advantage. Consistent with this rationale, carboxylate-modified nanoparticles showed significantly increased uptake in StcE-treated ileoids compared with untreated controls (Figure 6G). In contrast, uptake of neutral SM-102 lipid nanoparticles remained unchanged, while amine-modified nanoparticles displayed a slight, non-significant decrease in intracellular signal following glycocalyx disruption (Figure S7A, C). These findings support our proposed model of the glycocalyx as a charge-selective barrier regulating nanoparticle uptake (Figure 6H, S7B, S7D).

Collectively, these findings demonstrate that mucin glycoproteins constitute an important protective component of the intestinal ileal glycocalyx by limiting bacterial invasion and selectively restricting nanoparticle uptake according to surface charge. These results further highlight the utility of apical-out ileoids as a physiologically relevant platform for investigating interactions between the distal small intestinal epithelium and luminal microorganisms or nanomedicines.

## 4. Discussion

Conventional ECM-embedded organoids have limited access to the apical surface, whereas 2D monolayers do not fully preserve the three-dimensional architecture of the intestinal epithelium.^[10,64]^ In the present study, building on recent advances^[9,10,40]^, we established apical-out human ileoids in suspension culture, combining direct access to the apical surface with the 3D architecture of the organoid model. Although apical-out organoids have been used to study several aspects of intestinal physiology^[9,14]^, their application to nanomaterial research remains largely unexplored. Our findings demonstrate that human apical-out ileoids support quantitative measurement of endocytic nanoparticle uptake across an intact apical surface and, on that basis, allow the apical glycocalyx to be tested as a regulator of nanoparticle access to the epithelium.

Primary organoid models can exhibit donor- and passage-dependent variation, which is relevant when establishing standardized experimental conditions. In our analysis, donor-derived ileoids showed broadly similar global proteomic profiles, although donor 1 exhibited the greatest overall similarity to the available native ileal reference samples and was therefore selected for subsequent experiments. Given the limited number of native tissue reference samples, this finding should be interpreted as an indication of donor-specific differences rather than evidence that donor 1 represents the most physiologically representative line. Across serial passages, ileoids remained broadly stable, whereas ileoids maintained to passage 22 showed reduced size, increased circularity, altered expression of differentiation-associated markers such as ALPI and MUC2, and enrichment of proliferation- and senescence-associated pathways. These findings support the use of defined passage ranges for comparative experiments while indicating that the model remains relatively robust during extended culture. Beyond donor and passage variation, the ileoids retained several structural and functional characteristics of the native ileal epithelium. They displayed a well-developed brush border, apical glycocalyx, and intact cell-cell junctions, together with epithelial barrier function and evidence of absorptive activity. These features indicate that the apical-out configuration provides more than simply experimental access to the epithelial surface and retains key properties relevant to interactions with luminal cargo.

The model further expressed multiple endocytic pathways relevant to interactions with luminal cargo. Proteomic analysis demonstrated expression of components associated with clathrin-mediated and fast endophilin-mediated endocytosis, as well as caveolae-mediated and dynamin-independent pathways. Notably, apical-out ileoids expressed low but detectable levels of CAV1, the principal structural protein of caveolae, whereas CAV1 has been reported to be absent or poorly represented in the widely used Caco-2 intestinal epithelial model.^[65,66]^ Caco-2 cells remain the standard *in vitro* model for intestinal transport and cellular internalization.^[3,38,39,67]^ The caveolae-associated machinery in ileoids may therefore support uptake mechanisms that Caco-2 cells do not capture. Endocytic proteins localized to the apical membrane and cortical actin network as reported elsewhere, while CAV1 showed predominantly lateral-to-basolateral localization as in native intestinal tissue.^[68]^ Intracellular endocytic structures and fluid-phase HRP uptake provided functional evidence of active trafficking. Consistent with functional endocytosis at the apical surface, intact ileoids internalized all three nanoparticle types tested (Figure 6D-F), establishing that nanoparticle uptake can be quantified across an unperturbed human ileal epithelium, which is the prerequisite for the glycocalyx experiments below. These data establish the presence and activity of endocytic machinery rather than the relative contribution of individual pathways, which will require pathway-specific perturbation.

Whereas intracellular uptake machinery determines subsequent cellular internalization, nanoparticles must first interact with the extracellular epithelial interface. The direct accessibility of the apical epithelial surface in apical-out ileoids therefore provides an opportunity to investigate the intestinal glycocalyx directly. Studies in rodents have suggested that the intestinal glycocalyx forms a dense, mesh-like structure with pore sizes of approximately 30 nm, supporting a potential size-selective barrier function that restricts the access of luminal cargo to the epithelial surface.^[28]^ However, much of the current understanding of glycocalyx structure and function is derived from animal models, endothelial or immortalized cell systems, and the extent to which these observations reflect the human ileal epithelial surface remains incompletely understood.^[28,33,36]^

In the present study, we first established that apical-out human ileoids possess a distinct glycocalyx at the apical epithelial surface using complementary proteomic, fluorescence imaging, and ultrastructural approaches. Proteoglycans, CD44, and surface-attached mucins were detected at levels within about two-fold of ileal biopsies, WGA labelled a glycoconjugate-rich layer overlying the F-actin-positive surface, and electron microscopy resolved a defined extracellular layer over the microvilli. Together, these findings indicate that the apical surface of the ileoids is not simply an exposed epithelial membrane but retains a complex extracellular interface resembling the native intestinal glycocalyx. To investigate whether the mucin component of this glycocalyx contributes functionally to epithelial protection, we used the mucinase StcE to cleave surface-attached mucins from the apical surface.^[16,61]^ StcE treatment reduced UEA-1 staining at the apical membrane, without affecting cell viability, confirming effective cleavage while preserving the underlying epithelium. This allowed the mucin layer to be tested separately from other glycocalyx components. As a benchmark against published data, we first used the *S.* Tm wt*/*p*ssaG*-GFP reporter strain. Mucin cleavage increased intracellular GFP-positive foci, indicating enhanced bacterial invasion. This reproduces previous findings in jejunal enteroid and colonoid models for this *Salmonella* strain.^[16]^ The experiment therefore validates the model in a segment that is a major site of *Salmonella* colonization^[69]^.

Having established a protective role for the mucin-rich glycocalyx in limiting bacterial invasion, we next asked whether the same epithelial barrier also regulates interactions with non-biological luminal cargo. This is relevant to oral nanomedicine, where particle size, surface chemistry, and charge influence diffusion, adhesion, and uptake at the epithelial surface.^[33–36]^ We therefore used nanoparticles with distinct surface charge properties to investigate whether mucin cleavage altered their interaction with apical-out ileoids. Strikingly, mucin cleavage selectively increased uptake of negatively charged nanoparticles, whereas uptake of near-neutral SM-102 LNPs, which are compositionally equivalent to the LNPs used in approved mRNA vaccines but were formulated here without mRNA, was unchanged and positively charged nanoparticles showed a slight, non-significant reduction in intracellular signal. The selective enhancement of anionic nanoparticle uptake following mucin cleavage is consistent with a contribution of the glycocalyx to charge-dependent interactions at the epithelial surface.^[33,36]^ Removal of negatively charged mucin-associated components may reduce electrostatic repulsion between the nanoparticle surface and the epithelial interface, thereby increasing nanoparticle accessibility to the plasma membrane. However, the present experiments cannot distinguish between altered electrostatic interactions, changes in steric accessibility, increased particle adhesion, or secondary effects on cellular uptake pathways, which will require further investigation.Taken together, the glycocalyx is not a passive structural layer but a functional interface: it restricts bacterial access by physical exclusion, whereas its effect on nanoparticles is charge-dependent. Although charge-dependent interactions between nanoparticles and the intestinal glycocalyx have not, to our knowledge, previously been investigated in a human 3D apical-out ileal model, related charge-dependent interactions have been reported e.g. for the endothelial glycocalyx.^[33,36]^

Despite these advantages, several limitations should be considered when interpreting the physiological relevance of the model. Apical-out ileoids do not reproduce the full complexity of the intestinal microenvironment, including immune, stromal, vascular, neural, and microbial components that contribute to intestinal homeostasis and epithelial responses *in vivo*. Furthermore, although the apical-out configuration provides direct access to the epithelial surface, suspension culture does not reproduce physiological luminal flow or mechanical forces. The present model should therefore be considered complementary to, rather than a replacement for, more complex intestinal systems.

In conclusion, we established and characterized a reproducible apical-out human ileoid model that combines direct apical accessibility with the structural, molecular, and functional features of the human ileum. The model supports quantitative measurement of endocytic nanoparticle uptake across an intact epithelial surface, including of empty lipid nanoparticles whose composition matches that of approved mRNA vaccine formulations, and on that basis resolves the apical glycocalyx as a charge-selective regulator of nanoparticle access. That vaccine-like lipid nanoparticles are internalized by an intact human ileal epithelium is of particular interest for oral RNA delivery, a route that has so far been evaluated almost exclusively in cell lines; whether uptake of these empty particles translates into functional delivery of encapsulated mRNA is a question this platform is now well placed to address. More broadly, apical-out ileoids offer a tractable human system for studies of oral nanomedicine, epithelial transport, and host-microbe interactions at the ileal surface.

## Acknowledgements

This research was funded by the Swedish Research Council grant no. 2020-05186 (PA), Swedish Foundation for Strategic Research grant no. FFL18-0165 (MES) and the European Union’s Horizon 2021 research and innovation action (RIA) for GENEGUT (GA 101057491, PA and MH). RGP was supported by an Australian Research Council (ARC) Laureate Fellowship (FL210100107). H.M.G.B. acknowledges support from a Swedish Research Council Starting Grant (NT, 2022-04592) and funding from the European Research Council (ERC) under the European Union’s Horizon Europe research and innovation programme (Grant Agreement No. 101116289). The authors acknowledge the facilities and technical assistance of the Umeå Center for Electron Microscopy (UCEM) at the Chemical Biological Center (KBC), Umeå University, a part of the National Microscopy Infrastructure NMI (VR-RFI 2019-00217), and the facilities of the Microscopy Australia Research Facility at the Centre for Microscopy and Microanalysis at The University of Queensland. The BioVis platform of Uppsala University was used to conduct experiments using Leica Stellaris 5 confocal microscope.

## Author contributions

Conceptualization: D.S.V., M.E.S., P.A., and M.H.; data curation: D.S.V., A.H., F.T., J.E., M.L.D.M., W.F., and J.R.; formal analysis: D.S.V., A.H., F.T., J.E., and W.F., investigation: D.S.V., A.H., F.T., J.E., M.L.D.M., and W.F.; methodology: D.S.V., A.H., F.T., J.E., A.C.C.L., and M.L.D.M.; project administration: D.S.V., K.B., J.H., M.E.S., P.A., and M.H.; software: D.S.V., A.H., F.T., and J.E.; validation: D.S.V., A.H., F.T., J.E., M.L.D.M., W.F., M.E.S., P.A., and M.H.; visualization: D.S.V., F.T., and M.H.; supervision: M.E.S., P.A., and M.H.; resources: A.C.C.L., K.B., J.V., R.G.P., H.M.G.B., M.E.S., P.A., and M.H.; funding acquisition: R.G.P., H.M.G.B., M.E.S., P.A., and M.H.; writing – original draft: D.S.V. and M.H.; writing – review & editing: all authors.

## Data availability statement

The mass spectrometry proteomics data have been deposited at ProteomeXchange Consortium via the PRIDE partner repository as dataset identifier PRIDE: PXD084272 and are publicly available as of the date of publication. The data that support the findings of this study are available from the corresponding author upon reasonable request.

