## Supplementary figure for "Apical-Out Ileal Organoids: A Versatile Platform for Probing Gut-Nanomaterial Interactions"

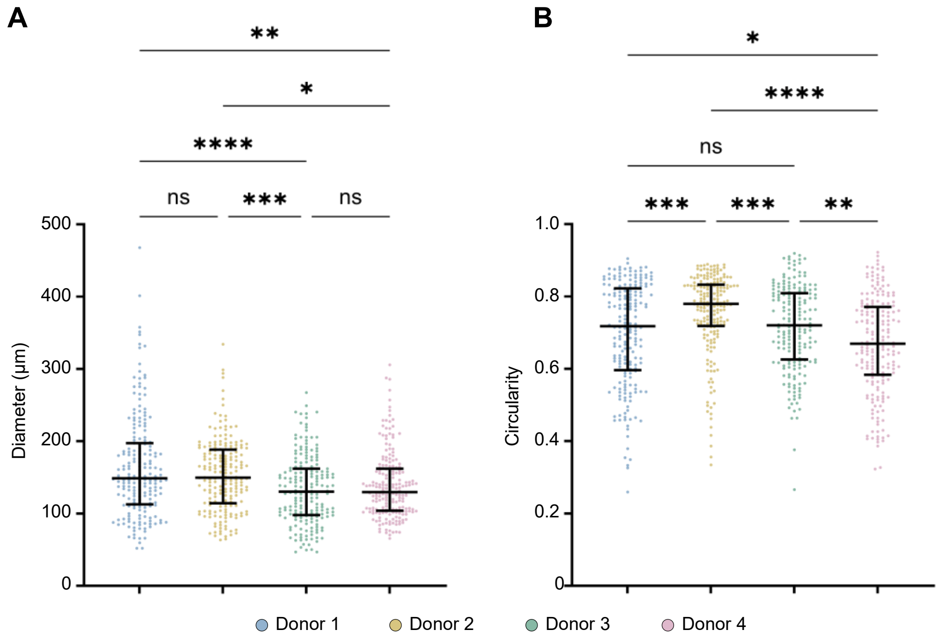


***Supplementary figure S1. Morphological characterization of apical-out ileoids from different donors. (A-B)*** *Quantification of* ***(A)*** *Feret's diameter and* ***(B)*** *circularity of ileoids derived from different donors. Data are shown as median and interquartile range (n = 200). P values were calculated by Kruskal-Wallis test; *p < 0.05, **p < 0.01, ***p < 0.001, and ****p < 0.0001.*

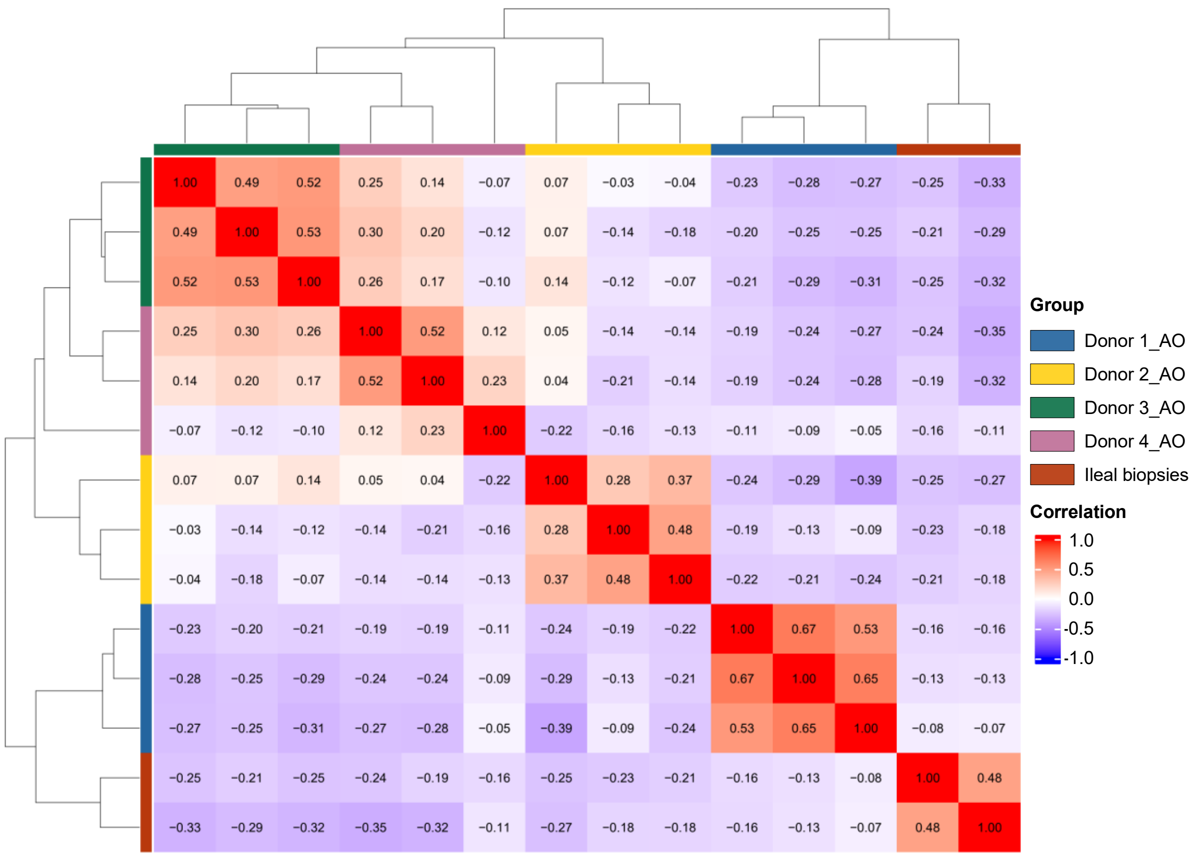


***Supplementary figure S2. Correlation matrix of proteomes from donor-derived apical-out ileoids and ileal biopsies.*** *Spearman correlation coefficients were calculated using vsn-normalized protein intensity values quantified across all sample groups. Heatmap colors represent the magnitude of pairwise correlations, ranging from low (blue) to high (red) similarity.*

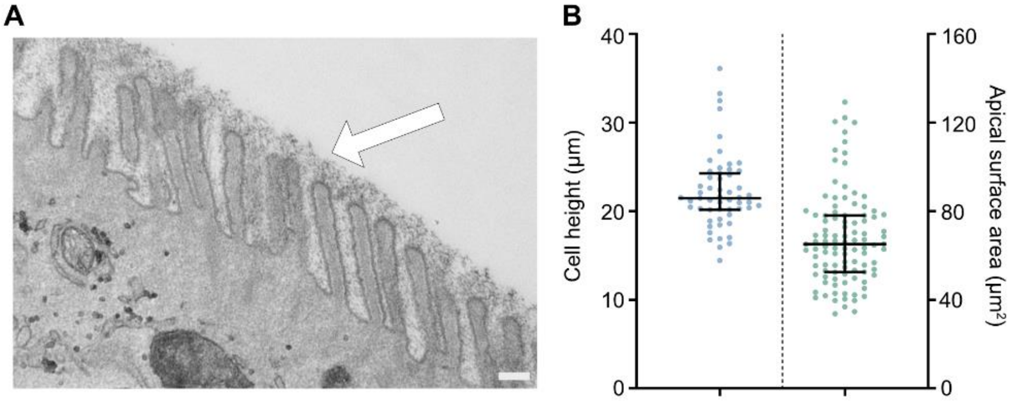


***Supplementary figure S3. Ultrastructural characterization of apical-out ileods. (A)*** *Representative TEM micrograph showing well-developed glycocalyx (white arrow). Scale bars, 200 nm.* ***(B)*** *Quantification of single-cell height and apical surface area in apical-out ileoids based on electron microscopy images. Data are shown as median and interquartile range (n = 55 for cell height measurement, n = 103 for apical surface area measurement). Donor 1 ileoids were used throughout.*

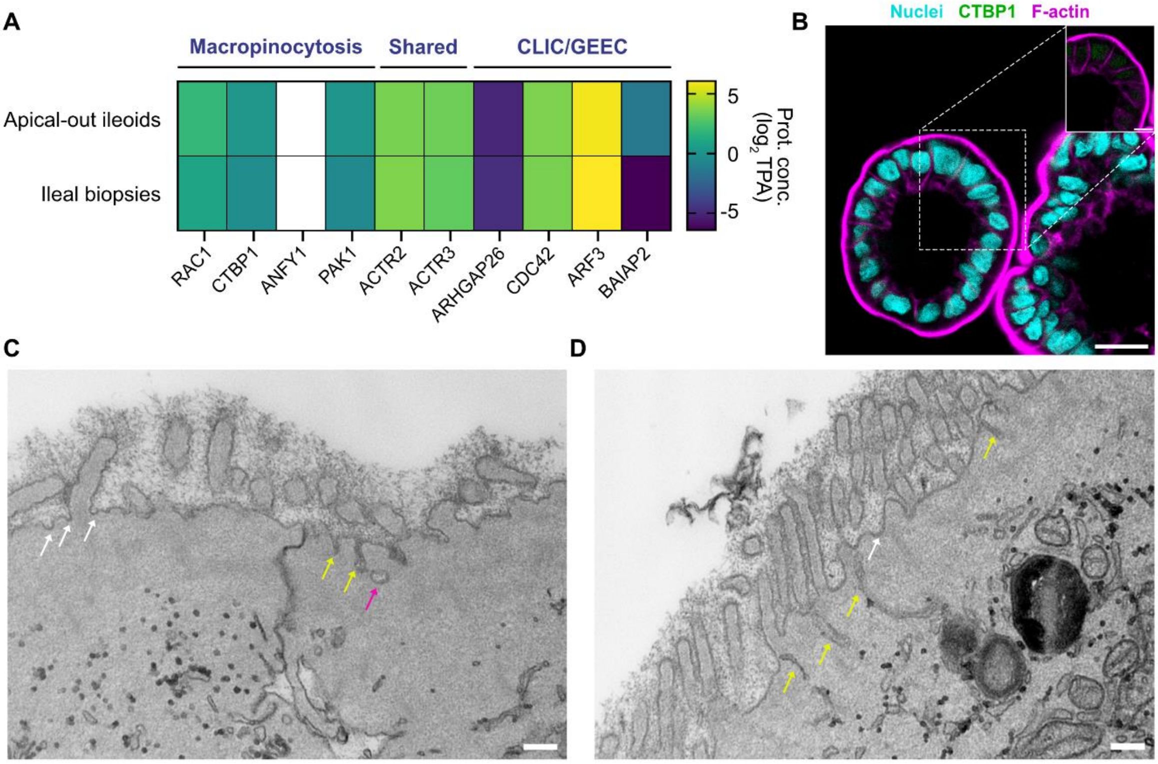


***Supplementary figure S4**. **Presence of dynamin-independent endocytosis in apical-out ileoids and ileal biopsies. (A)** Expression levels of representative protein markers of macropinocytosis and CLIC/GEEC endocytic pathways in apical-out ileoids compared with reference ileal biopsy samples. Data are shown as $log_2$ transformed of protein concentration after from total protein approach (TPA) quantification method. **(B)** Representative confocal images showing localization of CTBP1 protein involved in macropinocytosis. Scale bars, 20 µm; scale bars magnification, 10 µm. **(C and D)** Representative TEM images showing membrane pits (white arrows), narrow subapical tubular structures (yellow arrow) and intracellular ring-shaped membrane profiles (magenta arrow). The tubular and ring-shaped structures display morphologies consistent with endocytic intermediates associated with fast endophilin-mediated endocytosis (FEME) and/or CLIC/GEEC pathways. Scale bars, 200 nm. Donor 1 ileoids were used throughout.*

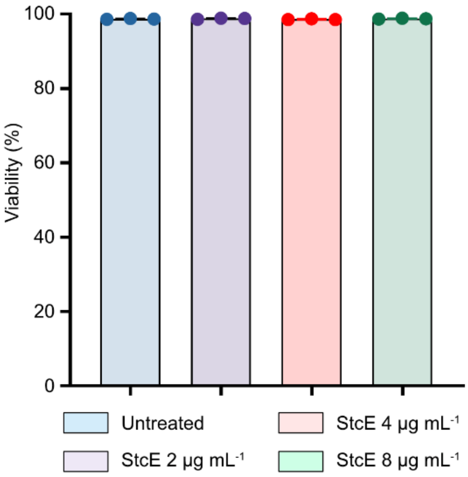


***Supplementary figure S5. Viability of apical-out ileoids following treatment with different concentrations of StcE.*** *Data are shown as mean ± SD (n = 3). Donor 1 ileoids were used throughout.*

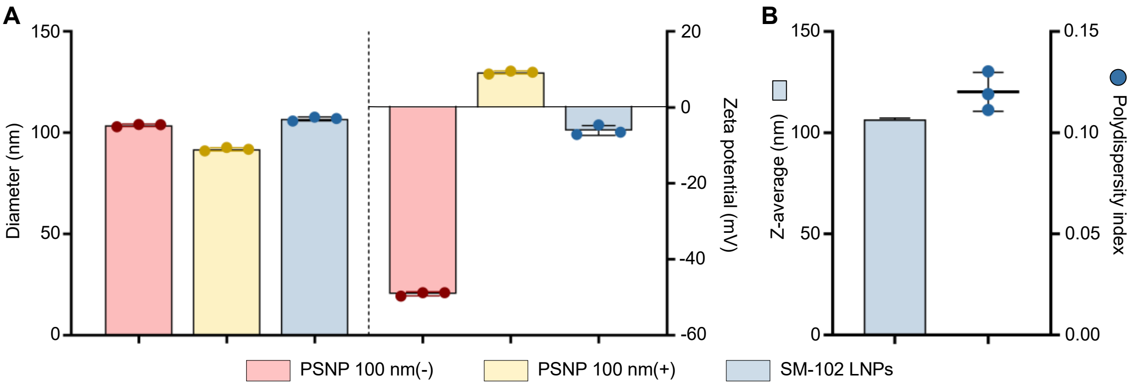


***Supplementary figure S6. Physicochemical properties of experimented nanoparticles.*** ***(A)*** *Hydrodynamic diameter and surface charge of formulated NPs measured by Nanoparticle Tracking Analysis (NTA). Presented results are reported as the median derived from the number-weighted distribution. Data are shown as mean ± SD (n = 3).* ***(B)*** *Z-average and polydispersity of formulated LNPs measured by Dynamic Light Scattering (DLS). Data are shown as mean ± SD across 3 technical replicates. Donor 1 ileoids were used throughout.*

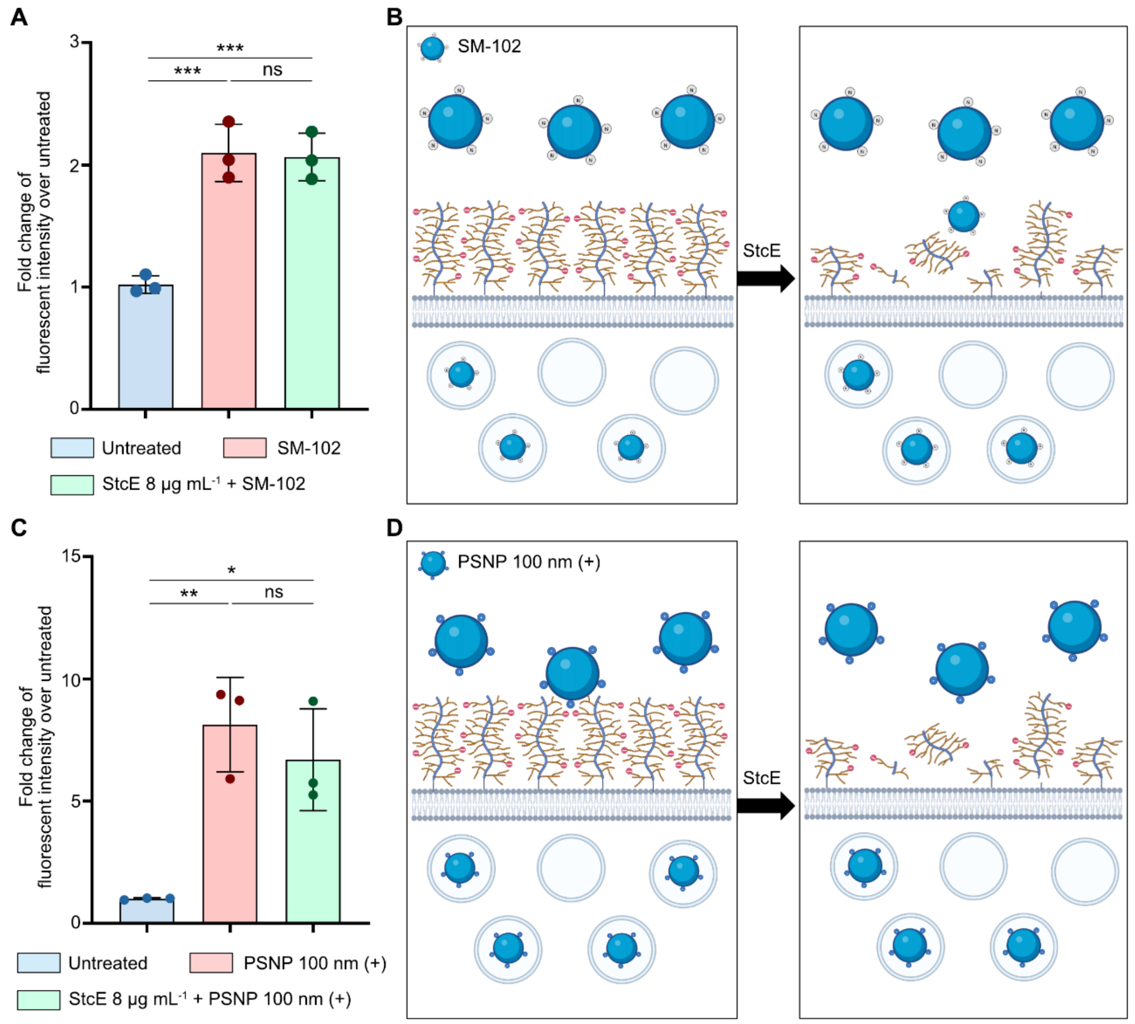


***Supplementary figure S7. Nanoparticle uptake in apical-out ileoids with and without StcE treatment. (A and C)*** *Quantification of intracellular nanoparticle fluorescence intensity in untreated and StcE-treated ileoids. The uptake of amine-modified polystyrene nanoparticles and SM-102 lipid nanoparticles was unchanged. Data are shown as mean ± SD (n = 3). P values were calculated by one-way ANOVA test; * p < 0.05, **p < 0.01, ***p < 0.001.* ***(B and D)*** *Proposed model illustrating the nanoparticle-glycocalyx interactions. Donor 1 ileoids were used throughout.*
